# Correction of frameshift mutations in the *atpB* gene by translational recoding in chloroplasts of *Oenothera* and tobacco

**DOI:** 10.1101/2020.09.11.293548

**Authors:** Irina Malinova, Arkadiusz Zupok, Amid Massouh, Mark Aurel Schöttler, Etienne H. Meyer, Liliya Yaneva-Roder, Witold Szymanski, Margit Rößner, Stephanie Ruf, Ralph Bock, Stephan Greiner

## Abstract

Translational recoding, also known as ribosomal frameshifting, is a process that causes ribosome slippage along the messenger RNA, thereby changing the amino acid sequence of the synthesized protein. Whether the chloroplast employs recoding, is unknown. I-iota, a plastome mutant of *Oenothera* (evening primrose), carries a single adenine insertion in an oligoA stretch of *atpB* (encoding a β-subunit of the ATP synthase). The mutation is expected to cause synthesis of a truncated, non-functional protein. We report that a full-length AtpB protein is detectable in I-iota leaves, suggesting operation of a recoding mechanism. To characterize the phenomenon, transplastomic tobacco lines were generated, in which the *atpB* reading frame was altered by insertions or deletions in the oligoA motif. We found that insertion of two adenines was more efficiently compensated than insertion of a single adenine, or deletion of one or two adenines. We further show that homopolymeric composition of the oligoA stretch is essential for recoding. Plants carrying a disrupted oligoA stretch have an albino-phenotype, indicating absence of indel correction. Our work provides evidence for the operation of translational recoding in chloroplasts. Recoding enables correction of frameshift mutations and can restore photoautotrophic growth in mutants that otherwise would be lethal.

## Introduction

Plastids (chloroplasts) are plant organelles that harbor their own genome (plastome) and are essential for many metabolic pathways, including photosynthesis and *de novo* synthesis of amino acids, nucleotides and fatty acids (Jarvis and López-Juez, 2013). As a result of endosymbiosis, they have evolved from a photosynthetic cyanobacterium that was engulfed by a eukaryotic cell (Herrmann, 1997; Bock, 2017). During subsequent coevolution of chloroplast and nuclear genomes, many plastid-encoded genes were lost or transferred to the nucleus (Timmis et al., 2004). Therefore, extant plastid genomes are small, and contain only coding information for approximately 120 genes in green plants (Bock, 2007; Barkan, 2011). The majority of these plastid-encoded genes are crucial for plant viability and required for photoautotrophic growth, including, for example, genes encoding the large subunit of RuBisCO, the reaction center subunits of the two photosystems, and subunits of the cytochrome *b_6_f* complex and the adenosine triphosphate (ATP) synthase. Other plastid-encoded gene products comprise components of the chloroplast gene expression machinery, such as the plastid-encoded bacterial-type RNA polymerase (PEP), the ribosomal RNAs (rRNAs), all transfer RNAs (tRNAs) and part of the ribosomal proteins. Due to its cyanobacterial ancestry, the chloroplast gene expression machinery is largely of bacterial type (Peled-Zehavi, 2007; Zoschke and Bock, 2018).

Precise genetic decoding is crucial to ensure the correct amino acid sequence in expressed proteins. During translation, spontaneous frameshifting is very rare with a frequency of about 10^-4^-10^-5^ per codon (Parker, 1989; Kurland, 1992). However, evidence has accumulated for the non-universality and plasticity of genetic decoding in many systems (Baranov et al., 2015; Atkins et al., 2016). Two major types of alternative genetic decoding can be distinguished: global codon reassignment and local transcriptional and translational recoding (Baranov et al., 2015). Global recoding leads to deviations from the standard genetic code. For example, in some bacteria, the stop codon UGA is reassigned to code for tryptophan (Yamao et al., 1985; McCutcheon et al., 2009) or glycine (Ivanova et al., 2014). Genetic code reassignment was also detected in plastids of the parasitic plant genus *Balanophora*, where UAG is reassigned from a stop codon to tryptophan (Su et al., 2019). At the local level, alternative genetic decoding can be caused by mRNA editing, transcriptional slippage or ribosomal frameshifting (Gesteland et al., 1992).

Transcriptional slippage or RNA polymerase stuttering is a process, by which the nascent transcript or transcriptional complex slips along the DNA template, resulting in the incorporation of additional or fewer nucleotides than encoded by the DNA template (Anikin et al., 2010). Transcriptional slippage mostly affects transcription of longer homopolymeric nucleotide tracts (Baranov et al., 2005), and results in the synthesis of a heterogeneous mRNA population. The slippage can occur during all phases of the transcription cycle (Anikin et al., 2010). RNA polymerase stuttering is frequently seen in some groups of RNA viruses, including paramyxoviruses (Jacques et al., 1994), the Ebola virus (Volchkov et al., 1995), the hepatitis C virus (Ratinier et al., 2008), and plants viruses of the *Potyviridae* family (Olspert et al., 2015). Here it often leads to the synthesis of more than one functional gene product from a single open reading frame (Volchkov et al., 1995; Hausmann et al., 1999). In a few cases, it also has been shown to compensate a frameshift mutation in the coding region (Baranov et al., 2005). For example, a single nucleotide deletion in the human apolipoprotein B (*apoB*) gene causes hypobetalipoproteinemia. However, the mutant allele is not a null allele, because, in addition to the expected truncated protein, some functional full-length ApoB protein was detected (Linton et al., 1992) that presumably was synthesized by transcriptional slippage.

Recoding at the level of translation can occur by programmed ribosomal frameshifting (PRF), a process in which the ribosome changes the frame it translates (Ketteler, 2012; Baranov et al., 2015; Atkins et al., 2016). PRF can occur in forward or reverse direction (relative to the 0-frame) as a consequence of the ribosome either skipping one nucleotide (+1) or slipping back one nucleotide (−1) (Ketteler, 2012). This can result in the production of two (or more) proteins from the same mRNA (Caliskan et al., 2017). Various signals are known to affect the efficiency of ribosomal frameshifting such as presence of a Shine-Dalgarno-like sequence upstream of the slip site and the mRNA secondary structure downstream of the slip site (Baranov et al., 2006). −1 frameshifting is more common than +1 frameshifting and, therefore, −1 PRFs has been analyzed in more detail (Belew and Dinman, 2015). −1 ribosomal frameshifting requires the slippery sequence motif X_XXY_YYZ, where X denotes any three identical nucleotides, Y denotes A or U, and Z is A, U, or C (Brierley et al., 1992). The P- and A-sites of the tRNA anticodon re-pair from XXY to XXX and YYZ to YYY, respectively (Jacks et al., 1988). In agreement with the consensus sequence, the slip site can be homopolymeric such as the A_AAA_AAA site in the adenomatous polyposis coli (APC) tumor suppressor protein of some nematodes (Baranov et al., 2011). The best-described bacterial −1 frameshift has the type A_AA.A_AA.G (with the codons of the original frame being separated by an underlined space, and the codons in the shifted frame by a dot) (Baranov et al., 2006). A pseudoknot in the mRNA structure is another motif that is often involved in successful ribosomal frameshifting. The pseudoknot consists of two or more stem-loop motifs and resides 3’ of the slippery site, separated from it by a spacer region of 5-9 nt (Ketteler, 2012). Together, slippery sequence and RNA secondary structures enable pausing of the ribosome and eventually frameshifting (Namy et al., 2006).

While the mechanisms of transcriptional slippage and translational frameshifting are distinct, both can occur on similar sites (Atkins et al., 2016). In *Escherichia coli*, synthesis of functionally distinct DNA polymerase III subunits (τ and γ) is achieved by ribosomal frameshifting, whereas in *Thermus thermophiles*, this is the result of transcriptional slippage followed by standard translation of the multiple mRNAs (Larsen et al., 2000). It is sometimes difficult to distinguish between low levels of transcriptional slippage and ribosomal frameshifting (Atkins et al., 2016). For example, recoding in the core protein of the hepatitis C virus was first ascribed to PRF, but later attributed to transcriptional slippage (Ratinier et al., 2008).

Ribosomal frameshifting in plants has been shown to occur on some viral RNAs (Miller and Giedroc, 2010; Atkins et al., 2016). Recently, it was reported that a <u>c</u>onserved <u>p</u>eptide <u>u</u>pstream <u>o</u>pen <u>r</u>eading frame (CPuORF) in the mRNA of *AT3G57170* is a possible candidate for ribosomal frameshifting in *Arabidopsis* (van der Horst et al., 2019). Furthermore, ribosomal frameshifting was detected *in vitro* using wheat-germ extract (Napthine et al., 2003). Finally, indirect evidence for PRF was reported in chloroplasts (Kohl and Bock, 2009). By generating transplastomic tobacco plants, it was shown that the bacterial IS150 transposon was mobilized inside the chloroplast, which results in the accumulation of transposition intermediates. Since the synthesis of IS150 transposase requires PRF (Polard et al., 1991; Vögele et al., 1991), PRF was proposed to exist in plastids.

Mutants in the chloroplast genome have been proven as a powerful tool in the analysis of chloroplast gene expression and photosynthesis (Schaffner et al., 1995; Landau et al., 2009; Greiner, 2012). Based on their chlorotic phenotypes, a collection of 51 spontaneous chloroplast mutants was isolated in evening primroses (*Oenothera*) (Kutzelnigg and Stubbe, 1974; Stubbe and Herrmann, 1982). Recently, this collection was systematically characterized by full plastome sequencing (Massouh et al., 2016). Chloroplast mutants affecting all photosynthetic complexes as well as the chloroplast translation machinery were identified. This collection also offers great potential to uncover novel mechanisms of chloroplast gene regulation (Greiner, 2012; Massouh et al., 2016). One of the most striking mutant phenotypes observed in this collection was the mottled phenotype of the I-iota mutant. It displays green spots on white leaves which grow with age, leading to a nearly green phenotype (Figure 1C in Massouh et al. 2016). The underlying mutation is a single adenine insertion (+1A) located within an oligoA stretch in the *atpB* gene, encoding the β-subunit of the plastid ATP synthase (Massouh et al., 2016). The AtpB protein is part of the α_3_β_3_ catalytic center harboring the nucleotide binding sites of the enzyme (Hahn et al., 2018). Here, we have analyzed the physiological and biochemical consequences of the +1A mutation in the I-iota mutant. We provide evidence for partial rescue of AtpB synthesis by ribosomal frameshifting. To characterize the recoding mechanism, a set of transplastomic tobacco mutants with mutated frameshift sites were generated and analyzed. Recoding efficiency was further assessed using a luciferase-based reporter system in *E. coli*. We present clear evidence for recoding in the chloroplasts depending on length and composition of the slip site and discuss its functional implications.

**Figure 1.**
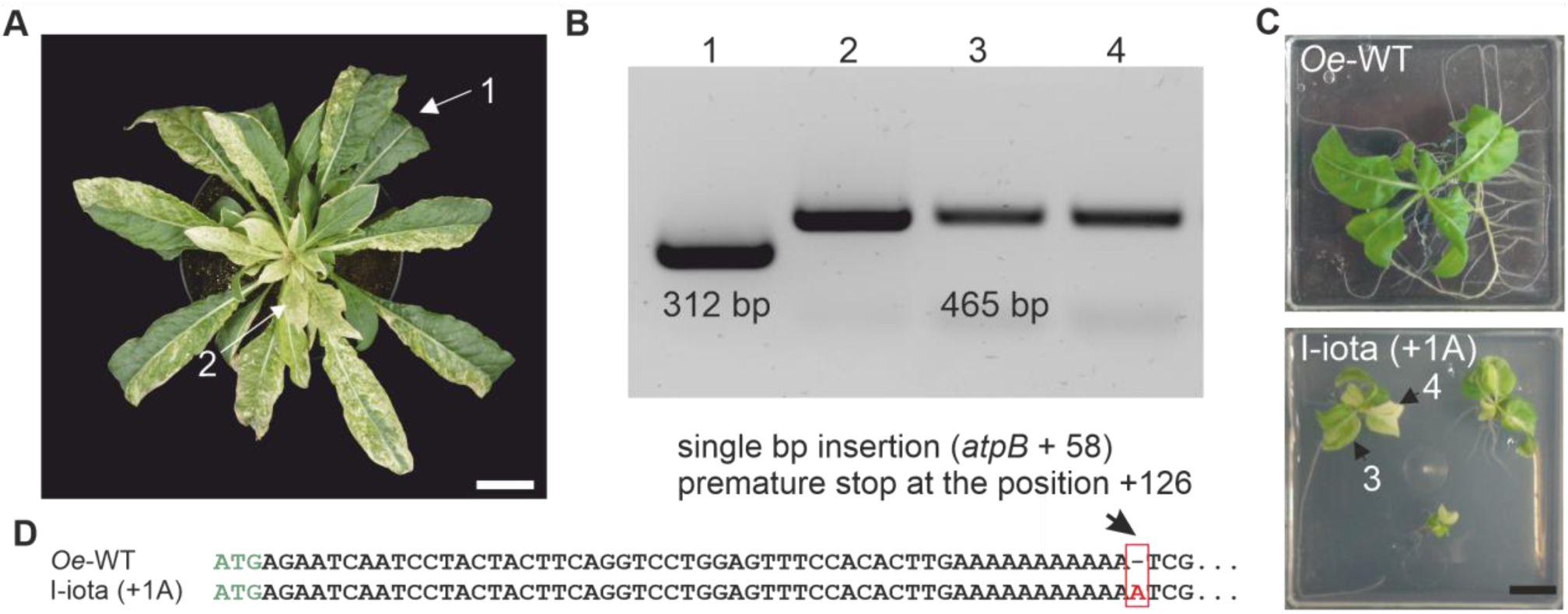
Phenotype and genotype of the I-iota mutant. **A**. Variegated *Oenothera* plant at the mature rosette stage (seven weeks old) carrying homoplasmic sectors containing the nursery plastome (green) [1] and homoplasmic sectors with I-iota chloroplasts (mottled) [2]. Scale bar: 5cm **B.** PCR confirming homoplasmy of mottled and green sectors (1: nursery plastome [312 bp], 2-4: I-iota plastome [465 bp]; note the absence of the 312 bp band inform lanes 2-4). **C.** Phenotype of wild type (*Oe*-WT) and I-iota homoplasmic plants cultivated under mixotrophic conditions in sterile culture. Lanes 3 and 4 in panel B confirm homoplamsy of the mottled plants. Scale bar: 1cm. **D**. 5’ part of the *atpB* gene of *Oe*-WT and I-iota (+1). The mutation at position +58 in an oligoA stretch of *atpB* in I-iota is framed in red. Green letters indicate the start codon.

## Results

### Molecular and biochemical characterization of the Oenothera I-iota mutant

I-iota was initially identified as chloroplast mutant displaying a mottled phenotype (Sears and Herrmann, 1985; Massouh et al., 2016). When the mutation is present in the homoplasmic stage, the plants cannot grow photoautotrophically. Therefore, to facilitate growth in soil, plants must be kept as variegated plants that are heteroplasmic and contain green nursery tissue (with wild-type chloroplasts) and pale sectors with mutated chloroplasts (Figure 1A). Taking advantage of biparental chloroplast inheritance in evening primrose, appropriate plants can easily be generated by a simple cross. To this end, a maintainer line with the chloroplast genome of *O. parviflora* is crossed to the I-iota mutant originating from *O. elata* (cf. Massouh et al. 2016; Greiner, 2012) (and Material and Methods). In order to confirm that the mosaic pattern of I-iota was not caused by incomplete vegetative sorting-out of the nursery plastome in the mutated I-iota sectors (Figure 1A) (cf. Greiner, 2012), tests for homoplasmy in green and mottled tissue were performed. The tests employed a length polymorphism in the plastid *clpP* gene that results in a larger PCR amplification product in the I-iota plastome of *O. elata* compared to the nursery plastome from *O. parviflora* (Greiner et al., 2008). A single fragment of 312 bp was detected in green parts of the plant, product of 465 bp was identified in mottled areas (Figure 1A,B), thus confirming the homoplasmic state of the I-iota mutation in the mottled leaf sectors. To further characterize the phenotype of the I-iota mutant, homoplasmic I-iota plants were grown under mixotrophic conditions in sterile culture with sugar supplementation. Compared to wild-type plants, homoplasmic I-iota mutants were strongly reduced in growth and displayed a pale or variegated phenotype (Figure 1C). Homoplasmy of the I-iota mutant in sterile culture was verified by PCR using the same marker as described above and confirmed for both the pale and green parts of the leaves (Figure 1B, lanes 3 and 4).

Even without the knowledge of the underlying mutation, a defect in the chloroplast ATP synthase was predicted for I-iota. Moreover, the AtpB subunit was correctly identified as targeted by the mutation (Herrmann et al., 1980; Sears and Herrmann, 1985). Indeed, the MS analysis of our work confirmed reduced amounts of AtpA, AtpB and AtpE in I-iota tissue compared to the wild type (Figure 2A). However, in the study of Sears and Herrmann (1985) an unexpected high molecular band was recognized, using their anti-AtpE antibody. In the present western blot analysis, no AtpE was detected in I-iota young leaves, but distinct AtpE bands with the expected size of 14.5 kDa were identified in intermediate and mature leaves (Figure 2B). No unspecific high molecular band was detected in I-iota tissue (Supplemental Figure 1), indicating the absence of an AtpB/AtpE fusion product as suggested by Sears and Herrmann (1985). Recently, a single adenine insertion (+1A) in *atpB* was identified as the only mutation in the I-iota plastome (Massouh et al. 2016). The mutation is located within an oligoA stretch near the 5’ end of the gene (Figure 1D). The identified mutation results in a premature stop codon and formation of a truncated protein (41 amino acids long; Figure 1D, Table 1). Confirming the previous report of Sears and Hermann (1985), the full-length AtpB protein was detected in I-iota (+1A) leaf tissue by immunoblot analysis (Figure 2B). Importantly, although the AtpB protein level was lower in I-iota (+1A) leaves than in the wild type (*Oe*-WT), no difference in the mobility of the protein band between the wild type and the I-iota was observed. Mature leaves of I-iota contained higher levels of AtpB than young leaves (Figure 2B), suggesting that functional AtpB accumulates over time. This observation can also explain the age-dependent greening phenotype of I-iota mutant plants (Massouh et al., 2016). Analysis of plastid ultrastructure by transmission electron microscopy confirmed the age-dependent developmental gradient in thylakoid membrane formation of I-iota (Figure 2D-F). Although mature leaves of I-iota still have slightly disorganized thylakoid membranes, they are rather similar to wild-type thylakoids (Figure 2C,F).

**Figure 2.**
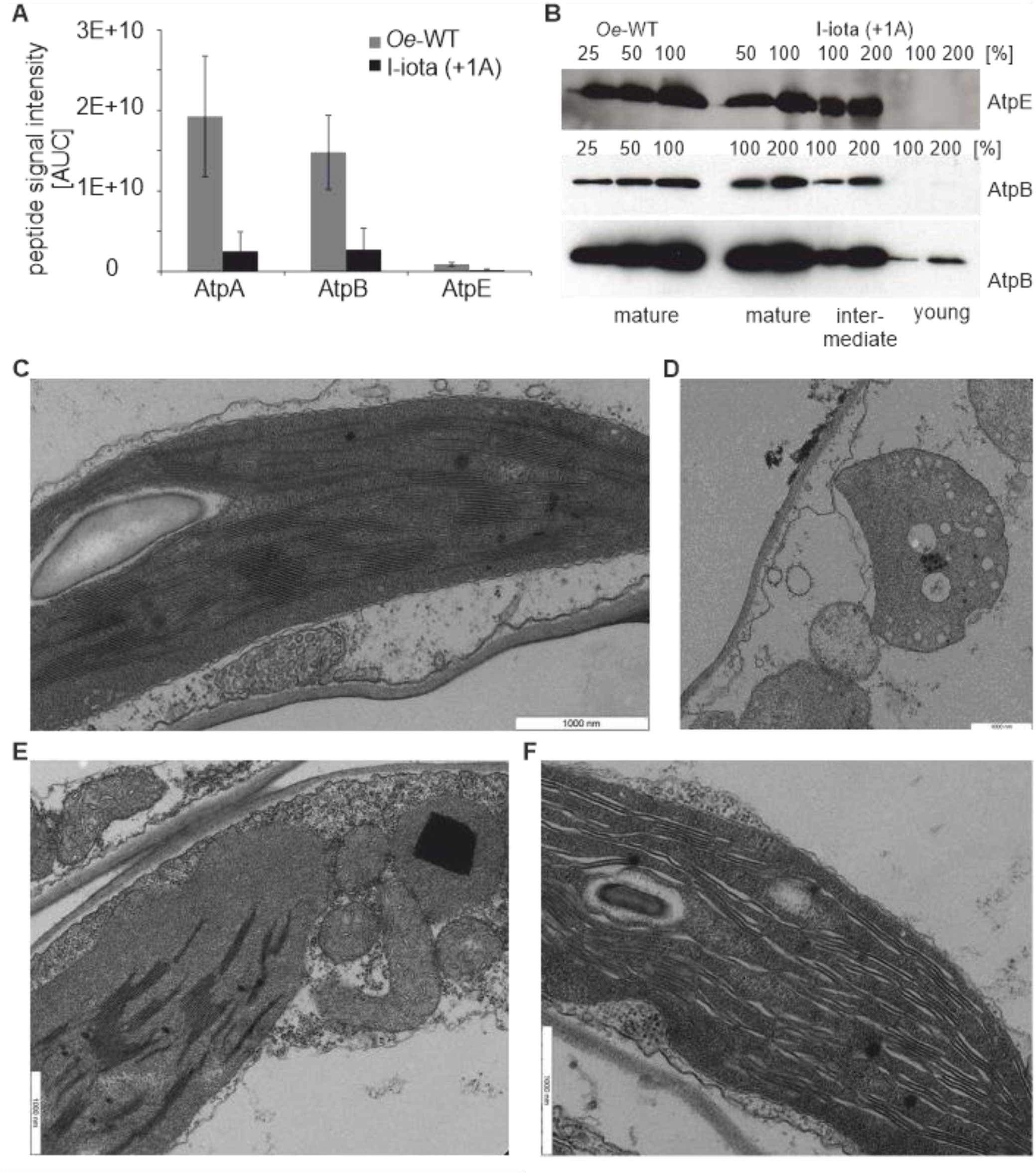
Analysis of ATP synthase and chloroplast ultrastructure in the I-iota mutant and the wild type. **A.** MS quantification of ATP synthase subunits in wild type and I-iota leaves (n=3). Signal intensities of peptides (AUC: area under the curve) of analyzed protein subunits, were normalized to BSA, which was used as internal control. **B.** Immunoblot detection of AtpE (14.5 kDa) and AtpB (ca. 53 kDa) in thylakoids from the wild type (*Oe*-WT) and I-iota leaves. Thylakoids were isolated from leaves of 6-8 week-old plants. For the I-iota mutant, leaf tissues of different ages were analyzed (young: top leaves of the rosette, intermediate: mottled leaves from the middle of the rosette, mature: greenish part of leaves from the base of the rosette; see Fig. 1A). Isolated thylakoids were tested for homoplasmy using the *clpP* marker prior to further analysis (see Fig. 1B). Thylakoids were loaded on the basis of the chlorophyll content, with 100% corresponding to 7 µg chlorophyll. The bottom panel (AtpB) shows a longer exposure of the same blot. **C-F**. TEM analysis of wild type [*Oe-*WT] and I-iota leaf tissue. **C**. Wild-type chloroplast. **D-F**. I-iota chloroplasts. **D**. Proplastid-like chloroplast. **E**. Chloroplast with developing thylakoid membrane. **F**. Mature wild type-like chloroplast. 7-week-old plants were investigated.

**Table 1.**
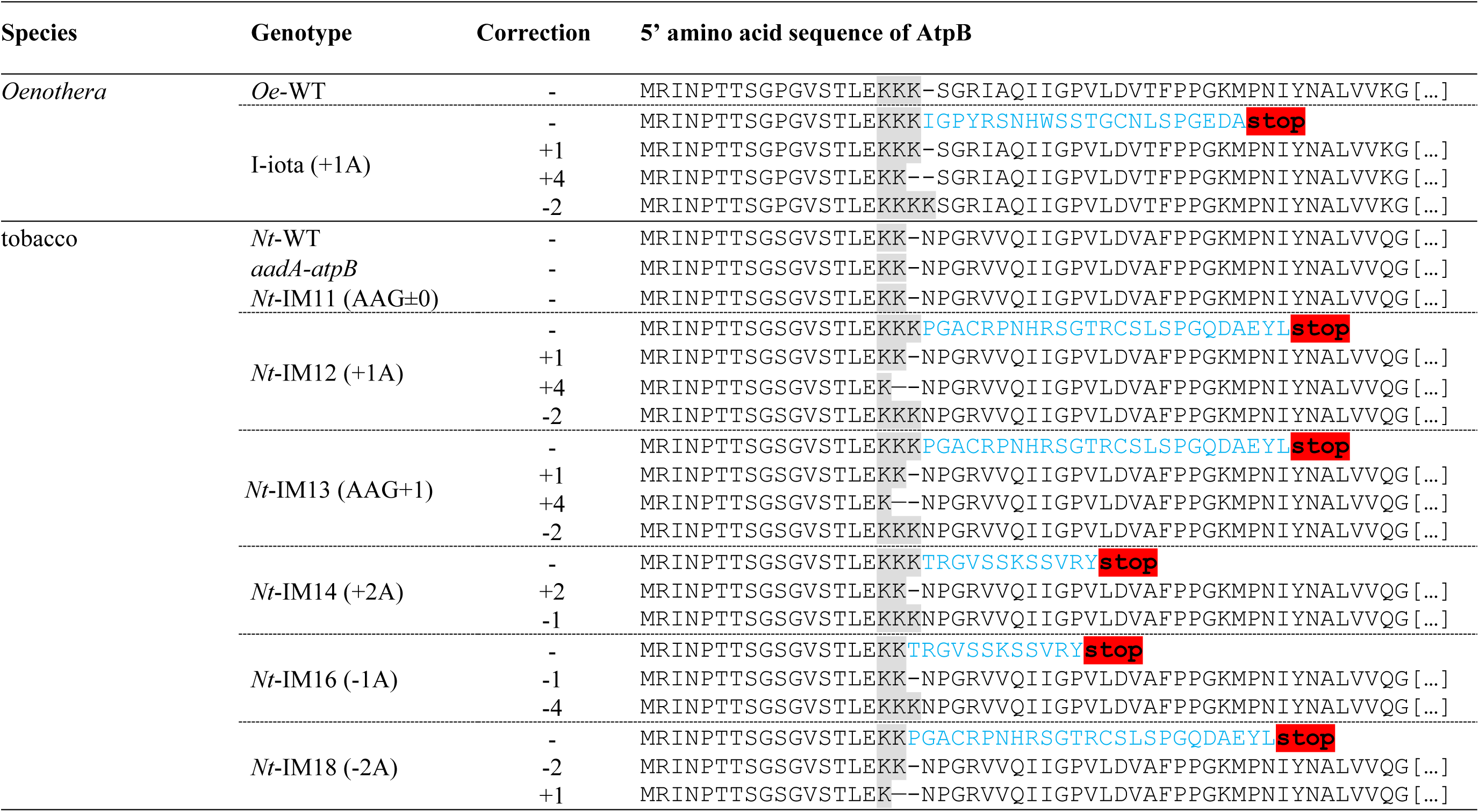
Possibility space of correction by recoding in Oenothera I-iota and tobacco transplastomic lines. The lysine stretch affected by the mutation is highlighted in grey. Blue letters indicate amino acids expected to deviate from the wild-type sequence according to the standard rules of genetic decoding. Premature stop codons are highlighted in red.

Presence of a full-length AtpB protein in I-iota tissue was verified by in-gel tryptic digestion followed by mass spectrometry (LC-MS/MS; Table 2). Finally, as *in vivo* measure of chloroplast ATP synthase activity, the proton conductivity of the thylakoid membrane (gH^+^) was assessed by decay kinetics of the electrochromic shift signal during a short interval of darkness. In mature leaves of I-iota, a residual ATP synthase activity of 29% compared to the wild type was detectable (Figure 3A), confirming that the chloroplast ATP synthase is indeed active in the I-iota mutant. As a consequence of the reduced ATP synthase activity, the entire photosynthetic apparatus of I-iota was affected. The chlorophyll content per leaf area was strongly reduced to 24% of the wild-type level, and also the chlorophyll *a/b* ratio was strongly decreased, indicating a substantial loss of photosystems (whose reaction centers exclusively bind chlorophyll *a*), while the antenna proteins (binding both chlorophyll *a* and *b*) were less severely affected (Table 3). Also, F_V_/F_M_, the maximum quantum efficiency of photosystem II (PSII) in the dark-adapted state, was clearly reduced in I-iota, pointing to PSII photoinhibition and the presence of uncoupled antenna proteins. Finally, light response curves of chlorophyll-*a* fluorescence parameters revealed that linear electron transport (calculated from PSII yield measurements) was strongly decreased in I-iota, and the acceptor side of PSII (here shown as the chlorophyll-*a* fluorescence parameter qL) became more rapidly reduced (Figure 3B). All these changes bear close similarity to photosynthetic defects previously reported for tobacco and mays (*Zea mays*) mutants with strongly diminished accumulation of ATP synthase (Rott et al., 2011; Zoschke et al., 2012), and can be ascribed to an increased proton motive force across the thylakoid membrane (because its consumption by ATP synthase is massively slowed down). This increased proton motive force leads to a strong acidification of the thylakoid lumen, which in turn slows down linear electron transport due to “photosynthetic control” of plastoquinol re-oxidation at the cytochrome b_6_f complex (Nishio and Whitmarsh, 1993), and also results in the stronger reduction of the PSII acceptor side, increased photoinhibition of PSII, and loss of photosystems (Rott et al., 2011; Schöttler et al., 2015). Taken together, these data strongly suggest partial compensation of the I-iota (+1A) mutation by a recoding mechanism that operates in I-iota tissue. Interestingly, analysis of the predicted structure of the *atpB* mRNA revealed presence of a potential slip site (oligoA stretch) and a stem-loop-type secondary structure downstream of the slip site (Figure 3C). Both motifs are known to be crucial elements for PRF. In contrast, a homopolymeric nucleotide sequence alone can be sufficient for induction of transcriptional slippage (see Introduction).

**Figure 3.**
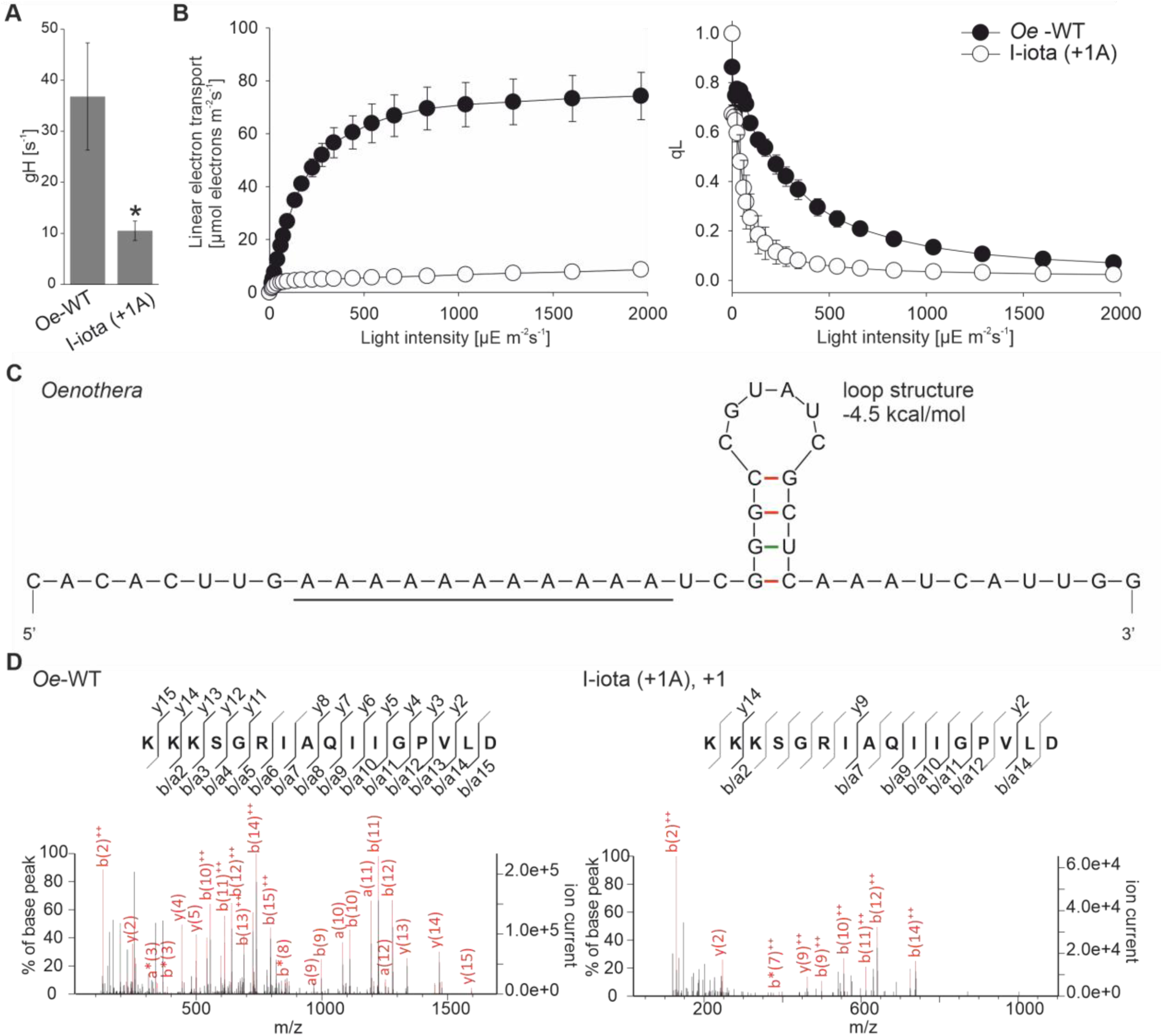
Photosynthetic parameters and analysis of target peptides in in the I-iota mutant and the wild type. **A.** ATP synthase activity determined from dark-interval decay kinetics of the proton motive force-related electrochromic shift signal. The asterisk indicates a significant difference compared to the wild type (t-test, *p* < 0.01, n=3-5). **B**. Light-response curves of linear electron flux (left panel) and chlorophyll-*a* fluorescence parameter qL (a measure of the redox state of the PSII acceptor side; right panel) in the wild type [*Oe*-Wt] and I-iota. 8 wild-type plants and 10 mutant plants were measured. Error bars represent the standard deviation. **C.** Predicted mRNA secondary structure of *atpB* near the slip site (oligoA stretch; underlined). **D.** LC-MS/MS fragmentation spectrum of the targeted AtpB peptide. The KKK stretch encoded at the slip site is identified as ‘b−’ and ‘y−’ type fragment ions.

**Table 2.** Identification of AtpB in *O*e*nothera* I-iota and transplastomic tobacco lines and the corresponding wild-type and control lines by in-gel tryptic digestion followed by LC MS/MS analysis. n.d. – not detected.

| Genotype | Nº of significant matches | Nº of significant sequences |
| --- | --- | --- |
| *Oe*-WT | 556 | 26 |
| I-iota (+1A) | 113 | 14 |
| *Nt*-WT | 249 | 34 |
| *aadA-atpB* | 77 | 24 |
| *Nt*-IM11 (AAG±0) | 155 | 31 |
| *Nt*-IM12 (+1) | 28 | 13 |
| *Nt*-IM13 (AAG+1) | n.d. | n.d. |
| *Nt*-IM14 (+2A) | 23 | 19 |
| *Nt*-IM16 (-1A) | 10 | 8 |
| *Nt*-IM18 (-2A) | 6 | 5 |

**Table 3.** Analysis of photosynthetic parameters. 3-5 plants were analyzed for both wild types (*Oe*-WT and *Nt*-WT), the I-iota (+1) mutant and the generated transplastomic tobacco lines. For each construct, at least two independent transplastomic lines were analyzed. Because no significant differences between these lines were observed, average data are shown for each genotype. Values are mean ± SD (n=3-5). Letters indicate samples that were not significantly different (*p* < 0.05) according to one-way ANOVA with Holm-Sidak post-hoc testing. ANOVA test was performed separately for greenhouse- and sterile culture-grown plants.

| Growth conditions | Genotype | Chlorophyll<br>[mg m<sup>-2</sup>] | Chlorophyll *a/b* | F<sub>V</sub>/F<sub>M</sub> |
| --- | --- | --- | --- | --- |
| Greenhouse | *Oe*-WT | 570.3 ± 12.8<sup>a</sup> | 3.6 ± 0.1<sup>a</sup> | 0.79 ± 0.01<sup>a</sup> |
| | I-iota (+1A) | 137.5 ± 51.2<sup>b</sup> | 2.7 ± 0.2<sup>b</sup> | 0.64 ± 0.02<sup>b</sup> |
| | *Nt*-WT | 438.4 ± 36.0<sup>c</sup> | 4.0 ± 0.2<sup>a</sup> | 0.83 ± 0.01<sup>a</sup> |
| | *aadA-atpB* | 414.7 ± 38.5<sup>c</sup> | 3.9 ± 0.1<sup>a</sup> | 0.81 ± 0.02<sup>a</sup> |
| | *Nt*-IM11(AAG) | 451.0 ± 23.5<sup>c</sup> | 3.8 ± 0.2<sup>a</sup> | 0.83 ± 0.01<sup>a</sup> |
| | *Nt*-IM14 (+2A) | 128.5 ± 17.0<sup>b</sup> | 2.6 ± 0.2<sup>b</sup> | 0.53 ± 0.08<sup>c</sup> |
| Sterile culture | *Nt*-WT | 297.3 ± 87.0<sup>ab</sup> | 3.50 ± 0.24<sup>a</sup> | 0.79 ± 0.04<sup>a</sup> |
| | *aadA-atpB* | 262.8 ± 47.9<sup>a</sup> | 3.42 ± 0.13<sup>a</sup> | 0.77 ± 0.03<sup>a</sup> |
| | *Nt*-IM11 (AAG) | 257.2 ± 63.1<sup>a</sup> | 3.50 ± 0.12<sup>a</sup> | 0.79 ± 0.03<sup>a</sup> |
| | *Nt*-IM12 (+1A) | 243.5 ± 70.1<sup>a</sup> | 2.34 ± 0.27<sup>b</sup> | 0.33 ± 0.14<sup>b</sup> |
| | *Nt*-IM14 (+2A) | 372.9 ± 78.1<sup>b</sup> | 2.78 ± 0.21<sup>c</sup> | 0.64 ± 0.18<sup>c</sup> |
| | *Nt*-IM16 (-1A) | 240.4 ± 95.7<sup>a</sup> | 2.17 ± 0.36<sup>b</sup> | 0.44 ± 0.23<sup>b</sup> |

Independent of the underlying molecular mechanism, mutation compensation is only partial and, therefore, will result in the formation of both in-frame and out-of-frame polypeptides (see Introduction). To characterize the process of mutation correction in I-iota at the protein level, we analyzed the lysine stretch encoded by the oligoA tract in which the mutation is located (Figure 1D). Since frame correction can occur in forward or reverse direction, different scenarios are possible (Table 1): (i) uncorrected gene expression will result in formation of a truncated polypeptide; (ii) +1 correction would lead to the synthesis of a protein identical to the wild type, and (iii) +4 or −2 correction would result in the synthesis of proteins that differ from the wild type in the number of lysines in the stretch (four or two lysines, respectively, instead of three in the wild-type protein). Consequently, resolving the number of lysines and the amino acid sequence downstream is crucial to the understanding of the correction mechanism. Unfortunately, standard trypsin digestion cannot be applied for mass spectrometric peptide identification, because trypsin cleaves at the carboxyl site of lysine and arginine residues (Tsiatsiani and Heck, 2015), and thus, is unsuitable to resolve the number of lysines in a peptide. As an alternative, we used in-gel digestion with glutamyl peptidase I (Glu-C) that cleaves the amino acid chain at aspartate and glutamate residues (Tsiatsiani and Heck, 2015). The peptide sequence information obtained from tandem mass spectrometry (MS/MS) was then analyzed against the *in silico* digested AtpB sequence from wild-type *Oenothera* and possible out-of-frame peptides that could be produced in the I-iota mutant (Table 1, Supplemental Table 1). No in-frame peptides corresponding to the expected truncated polypeptide were detected in I-iota samples, possibly suggesting that the truncated protein is unstable and condemned to rapid degradation. Instead, the same number of three lysines was detected in both the wild type and I-iota, revealing that the I-iota mutant indeed produces the wild-type AtpB proein and strongly suggesting that a +1 correction mechanism operates in the mutant chloroplasts (Figure 3D, Table 1).

### Reproduction of the I-iota mutation in transplastomic tobacco plants

Comparison of the plastid *atpB* sequences revealed that the oligoA stretch near the 5’ end of the coding region is conserved among higher plants and is also present in the cigarette tobacco (*Nicotiana tabacum*; Figure 4A). Moreover, a similar stem-loop structure was predicted to reside downstream of the oligoA tract in the *atpB* mRNAs of *Oenothera* and tobacco (Figure 3C, 4B,). We, therefore, attempted to reproduce the frame-correcting mechanism in tobacco chloroplasts, an experimental system amenable to an easy genetic manipulation. To obtain insights into the sequence requirements for reading frame correction, a set of transplastomic tobacco lines was generated (Figure 5A). To mimic the *Oenothera* I-iota mutation, a single adenine was inserted into the oligoA stretch within the 5’ part of the cloned tobacco *atpB* genes, generating plastid transformation vector pIM12 (+1A). The corresponding transplastomic lines will subsequently be referred to as *Nt*-IM12 (+1A) plants. To analyze the role of the oligoA sequence in the induction of recoding, a construct pIM13 (AAG+1A) was generated, in which the oligoA stretch was disrupted by replacing adenines with guanines in the third codon position (Figure 5A). These mutations do not alter the identity of the encoded amino acids, because both AAA and AAG triplets encode lysine. A similar construct with a disrupted oligoA stretch but without the frameshift mutation was generated as control [pIM11 (AAG); Figure 5A]. All transplastomic mutants were produced by biolistic chloroplast transformation using the *aadA* cassette as selectable marker that confers resistance to spectinomycin (Svab and Maliga, 1993; Figure 6A). Insertion of the *aadA* cassette alone did not affect plant growth and chloroplast ATP synthase activity (*aadA-atpB* control plants; Figure 5B; Rott et al., 2011).

**Figure 4.**
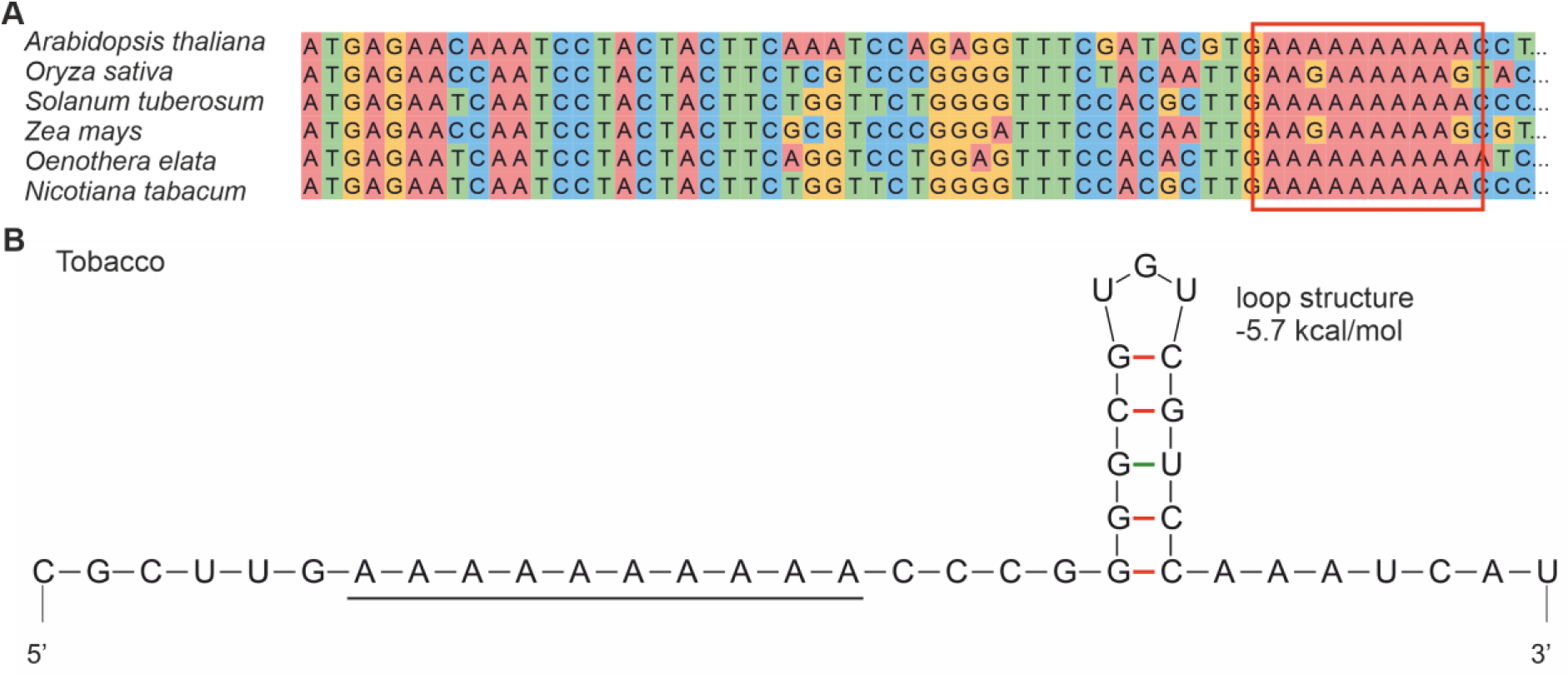
Sequence conservation and folding of the 5’ parts of the *atpB* coding region in different species. **A.** Alignment of the 5’part of the *atpB* coding region in higher plants. Note that all species have an oligopurine motif at the site where slippage occurs in *Oenothera*. the homopolymeric oligoA stretch (boxed in red) is conserved in *Arabidopsis thaliana*, *Solanum tuberosum*, *Oenothera elata*, and *Nicotiana tabacum*. **B.** Predicted mRNA secondary structure of *atpB* near the slip site (oligoA stretch; underlined) in tobacco.

**Figure 5.**
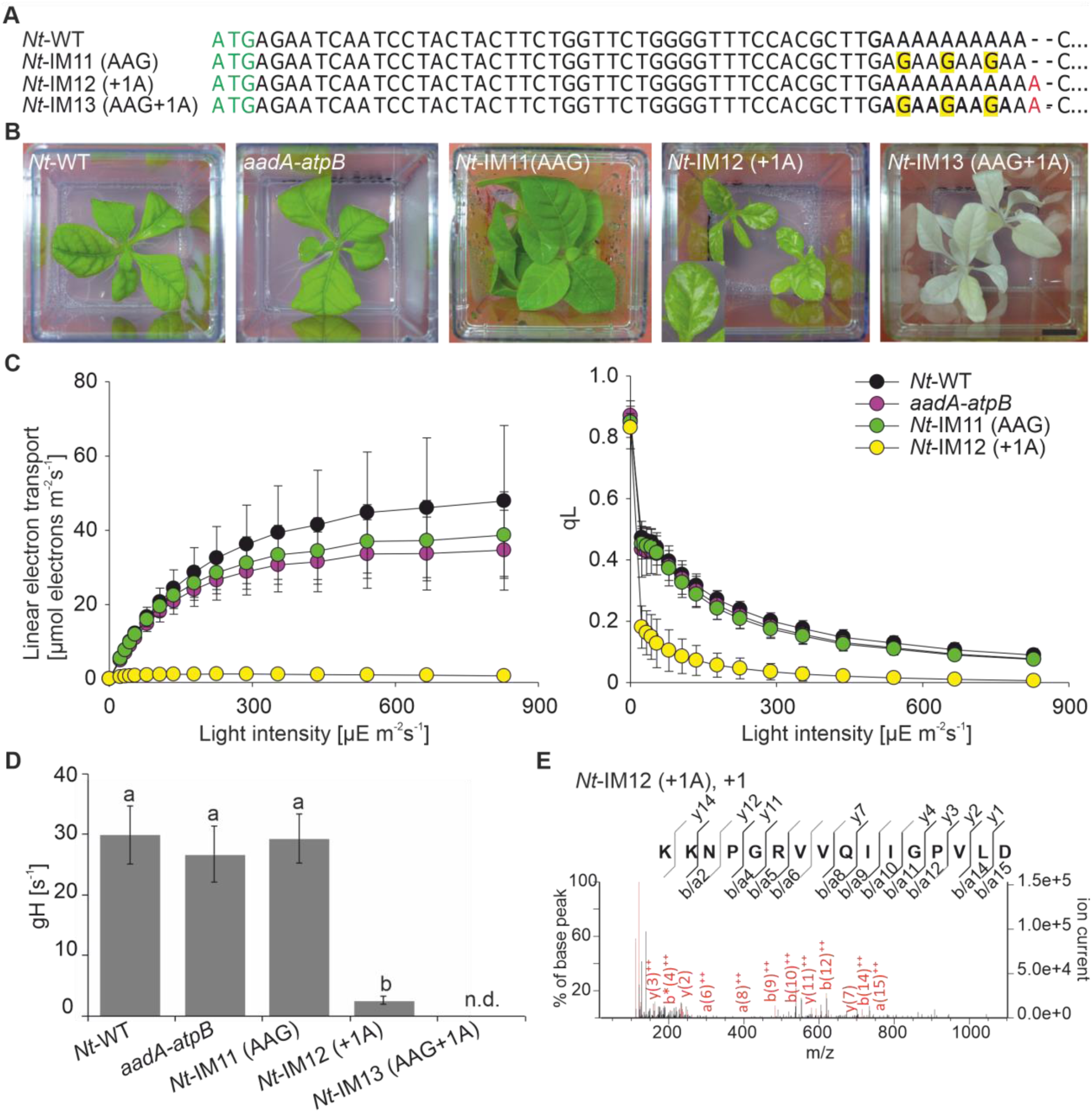
Generation and characterization of transplastomic tobacco plants carrying the I-iota mutation and/or disrupting the upstream oligoA stretch. **A.** Mutants generated to analyze recoding in tobacco. The sequence of the 5’ part of *atpB* from the start codon (green) to the oligoA stretch are presented. Red letters represent insertions. Adenines replaced by guanines are highlighted in yellow. **B.** Phenotypes of transplastomic tobacco lines grown in sterile culture on sucrose-containing medium. **C.** Light-response curves of linear electron flux (left panel) and chlorophyll-*a* fluorescence parameter qL (a measure for the redox state of the PSII acceptor side; right panel) in the wild type (*Nt*-Wt) and transplastomic tobacco lines. 10 wild type plants and 3-10 plants per transplastomic line were measured. For each construct, at least two independently generated transplastomic lines were analyzed. Error bars represent standard deviation. **D**. Activity of the ATP synthase. 10 wild type plants and 3-10 plants per transplastomic line were measured. For each construct, at least two independent transplastomic lines were analyzed. Columns display average data and standard deviation. Columns bearing the same letter indicate samples that were not significantly different (*p* < 0.05) according to one-way ANOVA with Holm-Sidak post-hoc testing. n.d – not detected. **E.** Analysis of the targeted AtpB peptide by LC-MS/MS (cf. Figure 3D).

**Figure 6.**
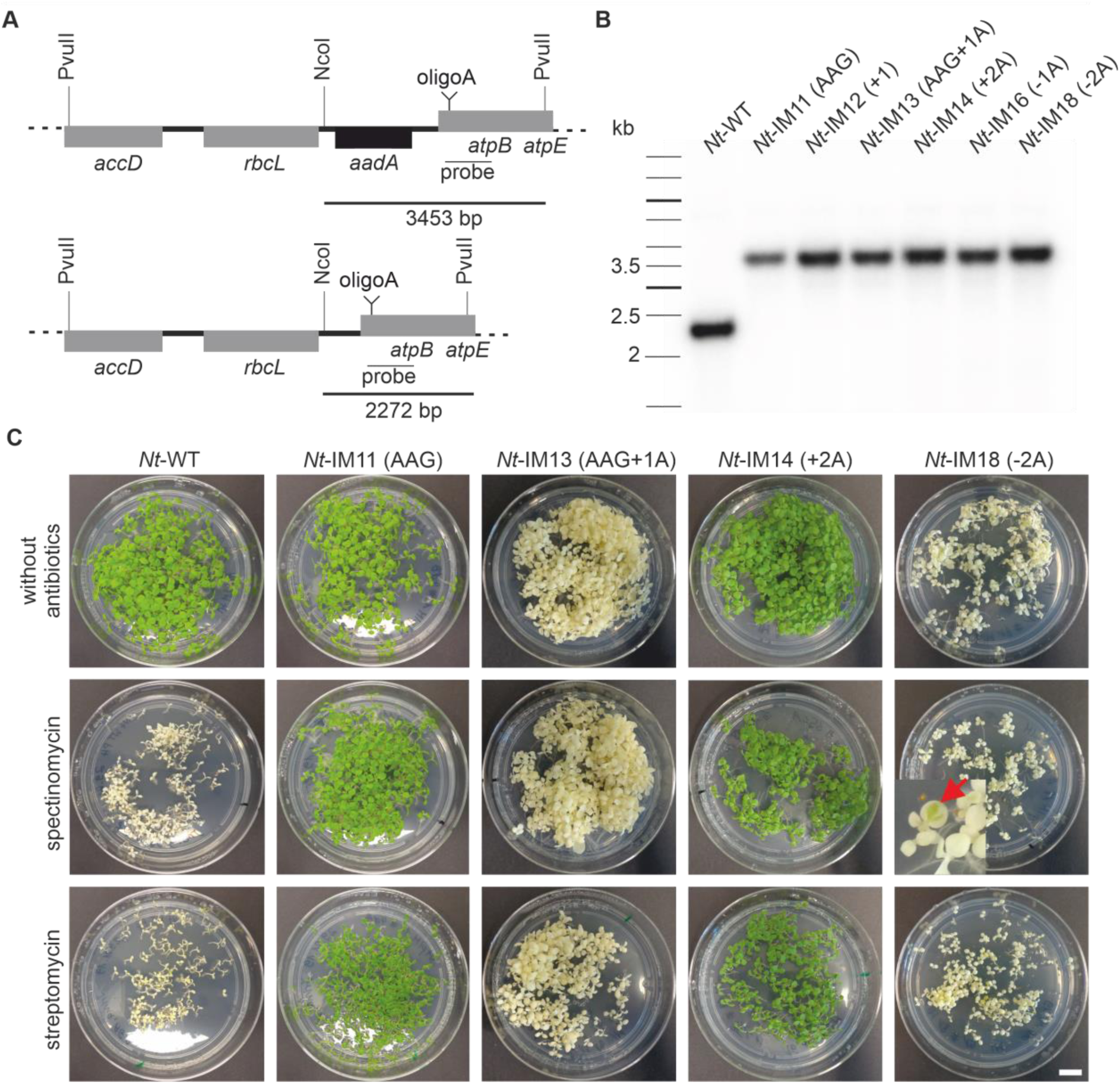
Confirmation of the homoplasmic state of transplastomic tobacco mutants. **A.** Physical maps of the plastome region of transplastomic lines (top) and wild type (bottom) (modified after Rott et al., 2011). Tobacco genes are shown as grey boxes, the black box represents the *aadA* cassette. Restriction sites used for RFLP analysis are indicated. **B.** Restriction fragment length polymorphism analysis of transplastomic plants. DNA was digested with PvuII and NcoI. The location of the hybridization probe is indicated in **A**. The wild type (*Nt*-WT) shows the expected restriction fragment of 2.3 kb, while all transformants show only the transplastomic fragment of 3.5 kb, confirming their homoplasmic state. **C**. Inheritance test. Seeds were germinated on synthetic medium without antibiotic (control) or medium supplemented with either streptomycin or spectinomycin. *Nt*-IM11 (AAG) and *Nt*-IM14 (+2A) seeds were derived from homoplasmic plants, *Nt*-IM13 (AAG+1) and *Nt*-IM18 (−2A) seeds were obtained from periclinal chimeras with mutated chloroplasts in their L2 layer (cf. Greiner, 2012) and Material and Methods). The red arrow indicates rare green spots on *Nt*-IM18 (−2A) mutant leaves. Scale bar: 1cm.

Replacement of three adenines with guanines in third codon position resulted in a wild-type-like phenotype [*Nt*-IM11 (AAG±0; Figure 5B], as expected. By contrast, *Nt-*IM12 (+1A) plants carrying the same mutation as I-iota (+1A) resembled the mottled phenotype of the *Oenothera* mutant (Figures 1A,C and 5B). Similar to *Oenothera* I-iota plants, transplastomic *Nt*-IM12 (+1A) plants were unable to grow autotrophically. Interestingly, combined disruption of the oligoA stretch and insertion of the I-iota mutation led to an albino phenotype. Transplastomic *Nt*-IM13 (AAG+1) plants are white and resemble the albino phenotypes of transplastomic *ΔatpB* knock-out mutants (Hager, 2002) and other loss-of-function mutants of the chloroplast ATP synthase (Karcher and Bock, 2002; Schmitz-Linneweber et al., 2005). This observation indicates that the oligoA stretch is of crucial importance for the frame-correcting mechanism.

Next, we analyzed photosynthesis in the transplastomic tobacco plants to assess their physiological similarity to the *Oenothera* I-iota mutant. No significant differences in linear electron flux and the redox state of the PSII acceptor side (qL) were detected between the wild type (*Nt*-WT), the *aadA-atpB* and the *Nt*-IM11 lines (Figure 5C). Interestingly, although chlorophyll content was similar in *Nt*-IM12 (+1A; the I-iota mutation) and the control lines (Table 3), all other parameters were reduced in *Nt*-IM12 (+1A). Linear electron flux was strongly decreased in *Nt-*IM12 (+1A), and the PSII acceptor side became reduced already in very low actinic light intensities (Figure 5C), similar to the *Oenothera* I-iota mutant. Also, F_V_/F_M_ was strongly reduced in *Nt*-IM12 (+1A), again pointing to PSII photoinhibition and the presence of uncoupled antenna proteins. This conclusion is further supported by the strongly decreased chlorophyll *a/b* ratio of *Nt*-IM12 (+1A), arguing for a predominant loss of the photosystems (Table 3). No analysis of photosynthetic performance was possible for *Nt*-IM13 (AAG+1), since the plants were completely white. In addition, activity of the ATP synthase was determined. As expected, no significant differences in ATP synthase activity between the wild type, the *aadA-atpB* control plants and *Nt*-IM11 (AAG) plants were detected (Figure 5D). By contrast, the line *Nt*-IM12 (+1A) displayed a strongly reduced activity of ATP synthase.

The observed phenotypes of the transplastomic tobacco lines suggested a varying degree of AtpB correction in the different lines. To explore the correlation between phenotype of the plants and synthesis of full-length AtpB protein, the accumulation of AtpB protein was analyzed by tryptic in-gel digestion followed by LC-MS/MS analysis. Full-length AtpB was detected in the wild type (*Nt*-WT), the control line *aadA-atpB*, and the *Nt*-IM11 (AAG)] and *Nt*-IM12 (+1A) mutants. No AtpB was detected in the white *Nt*-IM13 (AAG+1A) line (Table 2).

To compare the mechanism of reading frame correction with the I-iota mutant, the protein bands corresponding to AtpB were subjected to Glu-C digestion and analyzed by LC/MS-MS. As in I-iota, we identified identical peptides covering the slippery site in *Nt*-IM12 (+1A) and wild-type tissue (Figure 5E), suggesting operation of +1 correction also in the chloroplasts of tobacco (Table 1). No peptides derived from other reading frames could be detected (Table 1, Supplemental Table 1).

### Influence of mutations in the oligoA stretch on frameshift efficiency in E. coli

The phenotype of the *Oenothera* I-iota mutant and the transplastomic tobacco *Nt*-IM12 (+1A) line strongly suggests the presence of +1 frame correction in the chloroplasts of higher plants. Chloroplasts are of prokaryotic origin (Bock and Timmis, 2008) and have a bacterial-like translation machinery (Zoschke and Bock, 2018). Moreover, structure and subunit composition of CF_1_F_0_ resembles the ATP synthases of bacteria (Hahn et al., 2018). To test the possibility that the *atpB* reading frame correction also occurs in bacteria and, if it does, precisely measure its efficiency, a dual-luciferase reporter system in *E. coli* was employed (Kramer and Farabaugh, 2007). The dual-luciferase reporter assay is a powerful tool to analyze frameshifting efficiencies in various organisms and systems (Grentzmann et al., 1998; Kramer and Farabaugh, 2007; Ketteler, 2012). The assay is based on the Renilla luciferase serving as expression control, whereas the activity of the firefly luciferase depends on the expression of the test sequence (Grentzmann et al., 1998; Kramer and Farabaugh, 2007). Thus, the activity of the firefly luciferase relative to the Renilla luciferase reflects the translation efficiency (Grentzmann et al., 1998).

To determine the translation efficiency of our various *atpB* sequences, 150 nt of the 5’ part of the *atpB* gene from *Oenothera* and tobacco were cloned into the pEK4 plasmid (Kramer and Farabaugh, 2007) between the Renilla and firefly luciferase genes. In addition, several constructs varying in the length of the oligoA stretch or harboring a disrupted oligoA stretch (by placing guanines into the third codon position; see above) were generated (Supplemental Table 2). The translation efficiency measured in the constructs containing the *Oenothera* or tobacco wild-type sequence (pEK4-*Nt*-WT and pEK4-*Oe*-WT) was set as 100%. The pEK4-Nt-IM11 (AAG) construct, where the oligoA track of the tobacco WT sequence is destroyed by guanines, displayed an even enhanced efficiency compared to the wild-type sequences (Figure 7). By contrast, single adenine insertions caused a decrease in firefly luciferase activity relative to Renilla luciferase in *Oenothera* and the analogous tobacco constructs [pEK4-*Oe*-WT vs. pEK4-I-iota (+1) and pEK4-*Nt*-WT vs. pEK4-*Nt*-IM12 (+1A); Figure 7]. Similar to the *in planta* situation, disruption of the oligoA stretch in the single adenine insertion construct resulted in a significant reduction of the frameshift efficiency [pEK4-*Nt*-IM12 (+1A) vs. pEK4-*Nt*-IM13 (AAG+1A)]. Interestingly, +2A insertion in the oligoA stretch [pEK4-*Nt*-IM14 (+2A)] improved the translation efficiency by ca. 40%. Single or double adenine deletions [pEK4-*Nt*-IM16 (−1A) and pEK4-*Nt*-IM18 (−2A), respectively] did not exhibit effective frame correction and showed only approximately 5% of residual activity (Figure 7). In conclusion, also in the *E. coli*-based assay system, frame correction was detected in all constructs containing the single adenine insertion. Surprisingly, an even higher translation efficiency was detected with the construct with two adenines inserted.

**Figure 7.**
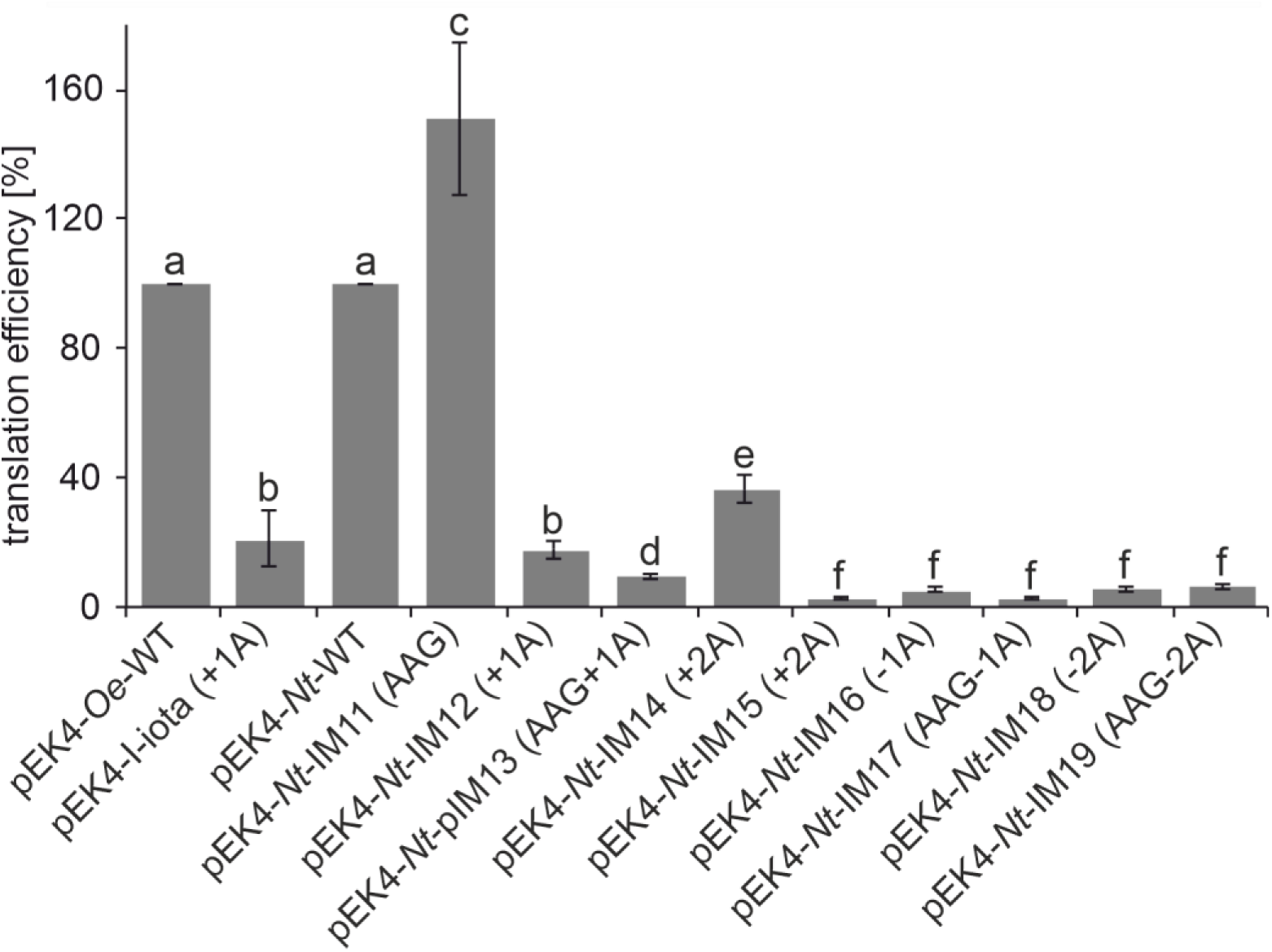
Impact of frameshift mutations in the oligoA stretch on translation efficiency as measured with the dual-luciferase system in *E. coli*. The error bars represent standard deviation from at least 3 independent biological replicates (i.e., bacterial colonies). Columns bearing the same letter indicate samples that were not significantly different (*p* < 0.05) according to one-way ANOVA with the Holm-Sidak post-hoc testing.

### Transplastomic tobacco plants with varying lengths of the oligoA stretch

The effective frame correction observed with the pEK4-*Nt*-IM14 (+2A) construct (Figure 7) raised the question how a +2A insertion into the oligoA stretch of *atpB* would affect the plant phenotype. To address this question, additional transplastomic tobacco lines were designed. Line *Nt*-IM14 (+2A) carries a +2A mutation, while lines *Nt*-IM16 (−1A) and *Nt*-IM18 (−2A) carry deletions of one or two adenines, respectively (Figure 8A). Homoplasmy of the generated transplastomic mutants was tested by restriction fragment length polymorphism analysis (Figure 6B), and presence of the mutations was verified by Sanger sequencing. Interestingly, *Nt-*IM14 (+2A) plants were indistinguishable from the wild type under mixotrophic conditions (Figure 8B). Moreover, although clearly retarded in growth, they were even able to grow photoautotrophically (Figure 9A). Compared to the wild type, *Nt*-IM14 (+2A) plants flowered 5-8 months later, but produced viable seeds that could be used for seed tests for homoplasmy (Figure 6C; cf. Material and Methods). Therefore, for *Nt*-IM14 (+2A) and the control lines *aadA-atpB* and *Nt*-IM11 (AAG) the full set of physiological parameters could be determined under both mixotrophic and autotrophic conditions (Figure 8C and 9C). Interestingly, plants with a single adenine deletion [*Nt*-IM16 (−1A)] displayed a similar phenotype as plants with a single adenine insertion [*Nt*-IM16 (−1A) vs. *Nt*-IM12 (+1A), Figure 8B vs. Figure 5B]. Both mutants exhibited pale green leaves with white areas. By contrast, the *Nt*-IM18 (−2A) line with the 2A deletion did not resemble the *Nt*-IM14 (+2A) phenotype. *Nt*-IM18 (−2A) plants were white with small green spots (Figure 8B), suggesting that the double adenine deletion cannot be efficiently compensated by recoding. Similar to *Nt*-IM13 (AAG+1), homoplasmic seeds could be obtained from stable periclinal chimeras (cf. Material and Methods). Seedlings that germinated in the presence or absence of spectinomycin were white with occasional green spots (Figure 6C). The visually observed phenotypes were further confirmed by the analysis of photosynthetic parameters: Nt-IM14 (+2) plants displayed a higher chlorophyll content compared to the other plant lines (Table 3), and the photosynthetic parameters of *Nt*-IM12 (+1A) and *Nt*-IM16 (−1A) were similar (Table 3, Figure 8C vs. 5C), suggesting comparable *atpB* correction efficiency. *Nt*-IM14 (+2A) displayed intermediate photosynthetic behavior when compared to WT and +1A or −1A plants (Figures 8C and 9C; Table 3).

**Figure 8.**
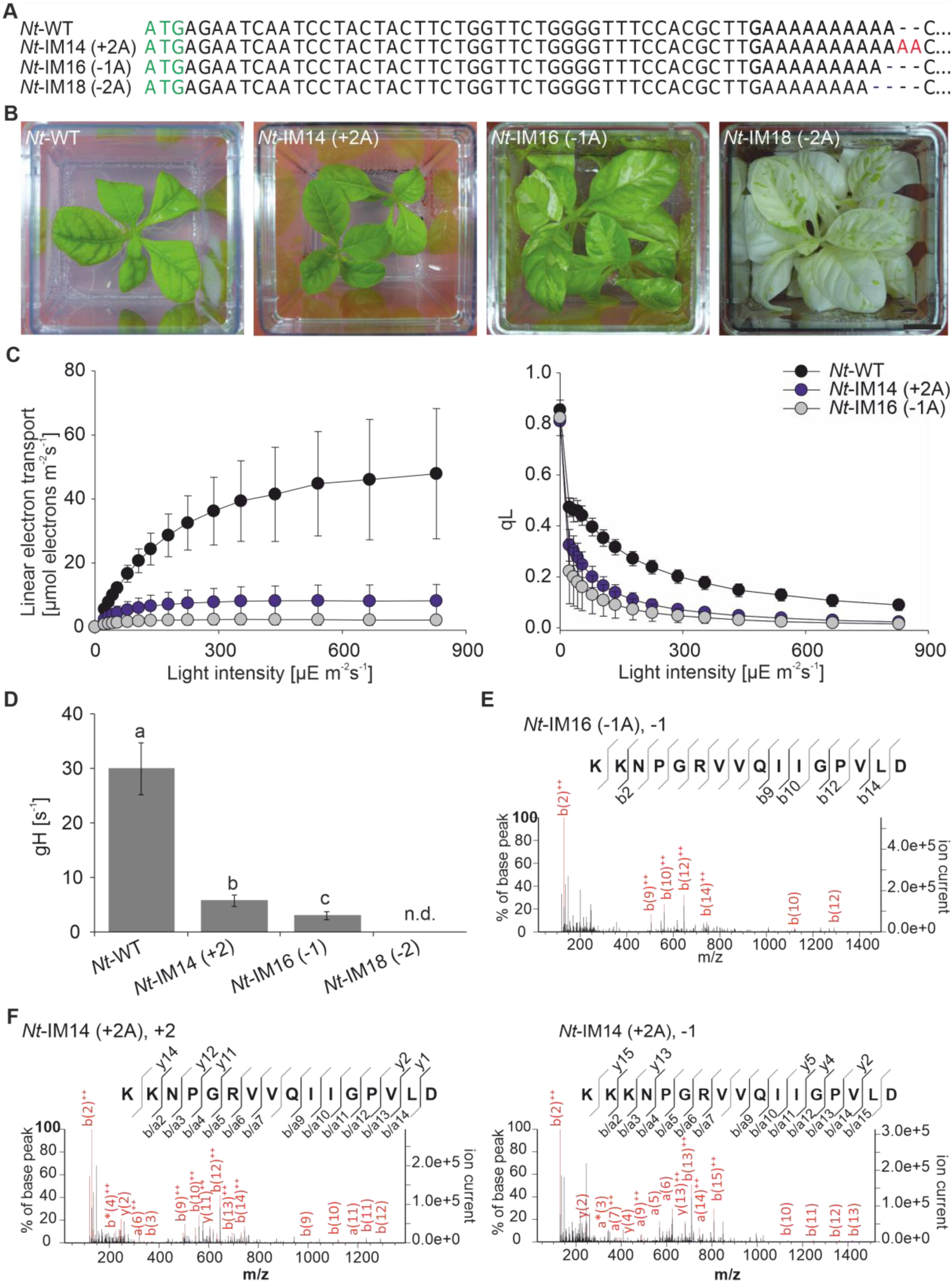
Generation and characterization of transplastomic tobacco mutants varying in the length of the oligoA stretch in *atpB*. **A.** Mutants generated to analyze recoding in tobacco. Red letters represent insertions, blue dashes denote deletions. **B**. Phenotypes of transplastomic tobacco lines grown in sterile culture. **C.** Light-response curves of linear electron flux (left panel) and the chlorophyll *a* fluorescence parameter qL (a measure for the redox state of the PSII acceptor side; right panel) in the wild type (*Nt*-Wt) and transplastomic tobacco lines. 10 wild type plants and 3-10 plants per transplastomic line were measured. For each construct, at least two independently generated transplastomic lines were analyzed. Error bars represent standard deviation. **D.** ATP synthase activity. 10 wild type plants and 3-10 plants per transplastomic line were measured. For each construct, at least two independent transplastomic lines were analyzed. Columns display average data and standard deviation. Columns bearing the same letter indicate samples that were not significantly different (*p* < 0.05) according to one-way ANOVA with the Holm-Sidak post-hoc testing. **E,F**. Analysis of peptides derived from the slippery site in AtpB by LC-MS/MS (for detail see Figure 3D).

**Figure 9.**
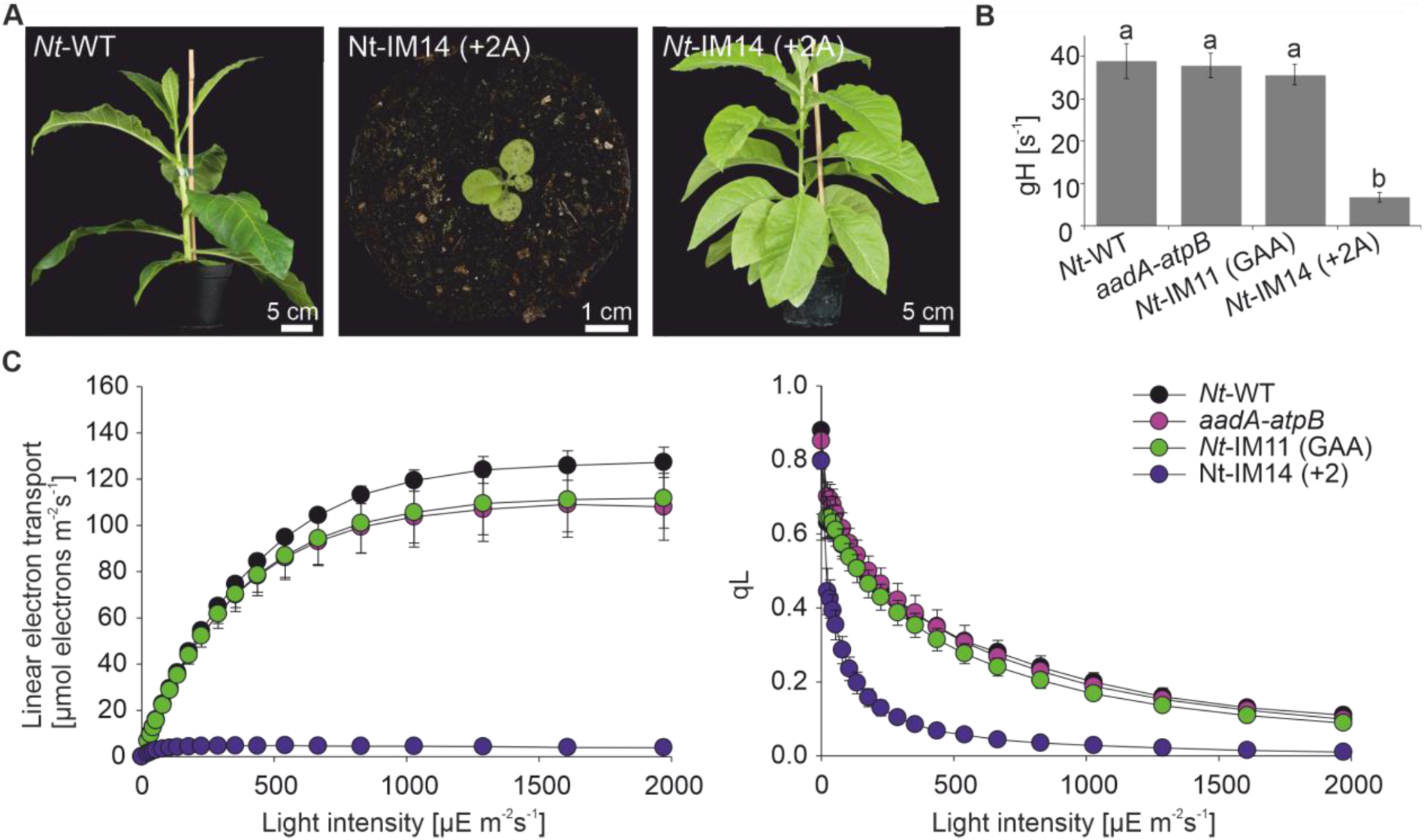
Correction of a +2 insertion in the *atpB* gene by recoding enables photoautotrophic growth. **A.** Growth phenotypes of wild type (*Nt-*Wt) and *Nt*-IM14 (+2A) plants cultivated in the greenhouse. Pictures were taken 7 weeks (left and middle) and 3 months (right photo) after germination. Note the strong retardation in growth of *Nt*-IM14 (+2A) mutants compared to the wild type. **B**. ATP synthase activity. *Nt*-WT, *aadA-atpB*, and *Nt*-IM11 (AAG) plants were 7 weeks old, *Nt*-IM14 (+2A) plants were 14 weeks old. 3 wild type plants and 3-5 plants per transplastomic line were measured. For each construct, at least two independent transplastomic lines were analyzed. Columns display average data and standard deviation. Columns bearing the same letter indicate samples that were not significantly different (*p* < 0.05) according to one-way ANOVA with Holm-Sidak post-hoc testing. **C**. Light-response curves of linear electron flux (left panel) and the chlorophyll *a* fluorescence parameter qL (a measure for the redox state of the PSII acceptor side; right panel) in the wild type and transplastomic tobacco lines. 4 wild type plants and 3-5 plants per transplastomic line were measured. For each construct, at least two independently generated transplastomic lines were analyzed. Error bars represent standard deviation.

Next, ATP synthase activity was assessed in the new set of transplastomic lines. The green *Nt*-IM14 (+2A) line displayed the highest ATP synthase activity among all mutants containing frameshift mutations (Figures 8D and 5D). In fact, when compared under mixotrophic and autotrophic growth conditions, *Nt*-IM14 (+2A) plants showed similar ATP synthase activity (19% and 17%, respectively; Figures 8D and 9B). By contrast, ATP synthase activity in *Nt*-IM16 (−1A) was at the detection limit, close to the decay kinetics of thylakoids with inactive ATP synthase. The electrochromic shift signal was not measurable in white *Nt*-IM18 (−2A) plants.

Finally, proteomic analysis of the transplastomic mutants was performed by tryptic in-gel digestion followed by LC-MS/MS. The AtpB protein was detected in all newly generated lines (Table 2). Peptides derived from the mutated site were generated by Glu-C digestion to gain insight into the correction mechanisms operating (Table 1). Peptides clarifying the amino acid sequence specified by the *atpB* mutations introduced into the transplastomic lines were identified in *Nt*-IM16 (−1A) and *Nt*-IM14 (+2A) (Figure 8E,F). Interestingly, both +2 and −1 corrections were detected in the Nt-IM14 (+2A) line, as evidenced by identification of peptides with two or three lysines (Figure 8F). A peptide resulting from +2 correction was detected in *Nt*-IM16 (−1A) (Figure 5E). No peptides were detected in the white line *Nt*-IM18 (−2A).

### Analysis of the atpB cDNA in Oenothera and transplastomic tobacco plants

Since RNA polymerase stuttering and ribosomal frameshifting can both occur at homopolymeric sequence motifs, it can be challenging to unambiguously distinguish between the two recoding mechanisms (Atkins et al., 2016). Transcriptional slippage generates a heterogeneous population of mRNAs that can be detected by cDNA analysis. Although seemingly straightforward, the high error rate exhibited by reverse transcriptases can complicate this approach (Roberts et al., 1988). To minimize this problem, we used a high-fidelity reverse transcriptase (a variant of Moloney murine leukemia virus reverse transcriptase, MMLV-RT, that was combined with a 3’-5’ exonuclease domain for proofreading). The enzyme exhibits an error rate that is three times lower than that of conventional reverse transcriptases (Arezi and Hogrefe, 2007). cDNA sequence analyses revealed that, in all transplastomic lines, the mutations introduced into the *atpB* locus were faithfully represented in the mature *atpB* mRNAs (Figure 10A). No sequence shifts or double peaks were detected in any of the chromatograms. A similar situation was observed for I-iota and the corresponding *Oenothera* wild type, although an overall somewhat more noisy sequence was obtained here for both the wild-type and the mutant (Figure 10B). To estimate the accuracy of cDNA synthesis and the sensitivity of the assay, we mimicked RNA polymerase stuttering by mixing total RNA from *Nt*-WT and *Nt*-IM14 (+2A) in different ratios followed by reverse transcription and sequence analysis of the amplified cDNA population (Figure 10C). When the amount of added *Nt*-WT RNA was ≥5%, an additional peak in the sequence chromatogram appeared (Figure 10C), revealing the presence of the heterogeneous *atpB* mRNA population. Absence of the double peak from the *atpB* sequences of the *Oenothera* I-iota mutant and the analogous transplastomic tobacco lines indicates that the level of RNA polymerase stuttering, if occurring at all, is below the 5% detection threshold. Given that the ATP synthase activity in the mutants reaches levels of more than 17% [in *Nt*-IM14 (+2A); Figures 8D and 9B], frameshift correction must occur mostly, if not exclusively, by ribosome frameshifting.

**Figure 10.**
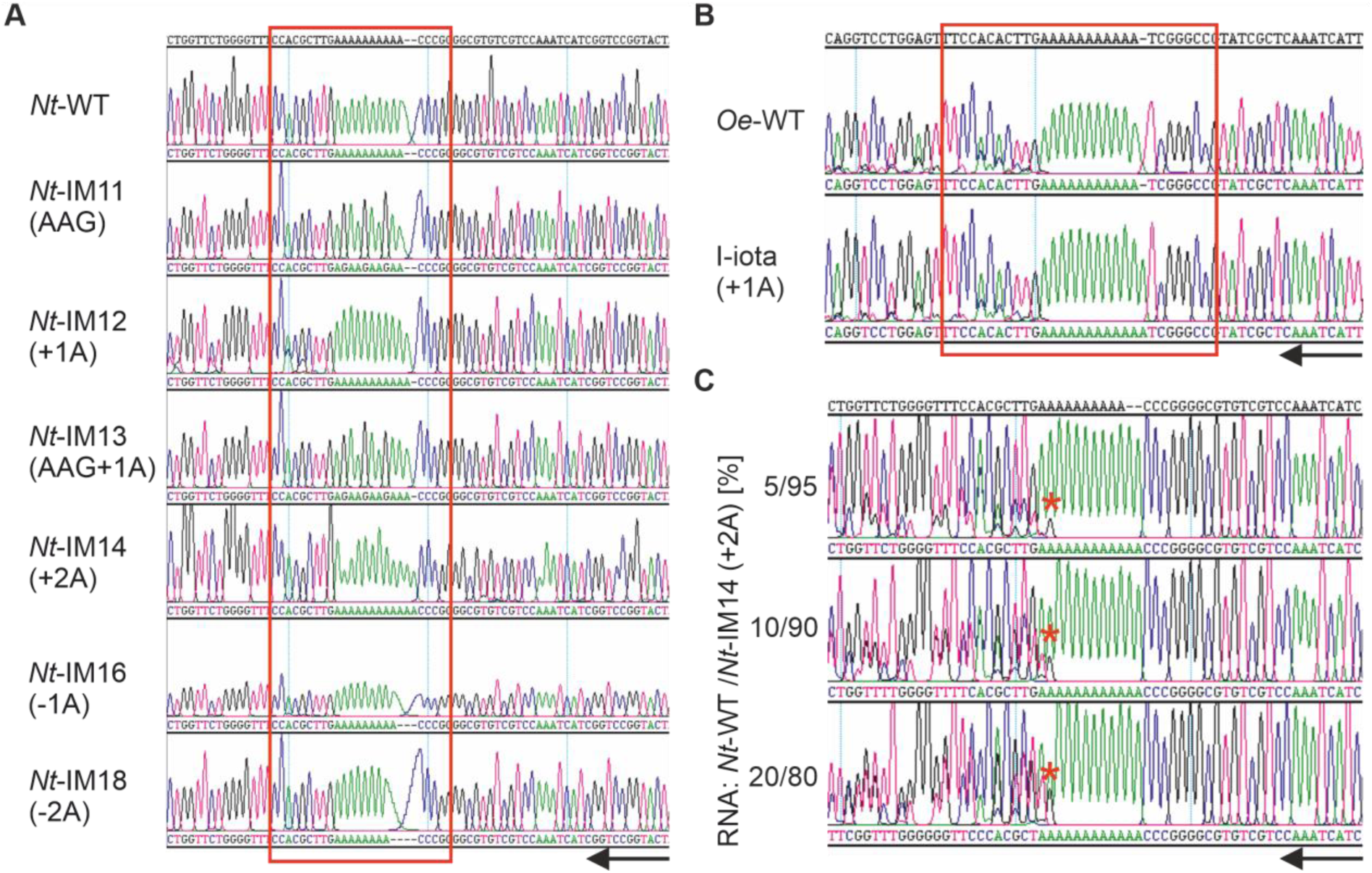
cDNA analysis to test for RNA polymerase stuttering in *atpB*. **A.** cDNA analysis of transplastomic tobacco plants and the wild type. The amplified cDNA population was sequenced, and typical Sanger sequencing chromatograms are presented. The sequence containing ther oligoA stretch is boxed in red. **B.** cDNA analysis of *Oenothera* wild type [*Oe*-WT) and I-iota (+1) plants. **C**. Sensitivity test for the detection of RNA polymerase stuttering. RNA was extracted from the wild type (*Nt*-WT) and the *Nt*-IM14 (+2A) transplastomic line, and the RNA samples were mixed in different ratios (indicated in %). cDNA was the synthesized, amplified and sequenced. Red asterisks indicate the additional peak, corresponding to the spiked-in wild-type RNA. Black arrows indicate the direction of the sequencing reaction.

## Discussion

Plastome mutants are an important tool for the analysis of non-Mendelian inheritance, the study of photosynthesis, and the analysis of the mechanisms of chloroplast gene expression (Greiner, 2012; Sobanski et al., 2019). Frameshift mutations usually lead to disruption of protein function, unless they are transcriptionally or translationally compensated (Ketteler, 2012; Atkins et al., 2016). Although such recoding is believed to exist in almost all organisms, only very few examples have been found in plants to date (see Introduction).

### Efficient recoding requires a homopolymeric sequence

Recoding based on transcriptional slippage relies on homopolymeric sequences (Baranov et al., 2005), while PRF can occur on either homopolymeric or heteropolymeric slip sites (Baranov et al., 2011). Transplastomic tobacco plants, where in addition to the frameshift mutation, adenines at the third codon position of the oligoA stretch had been replaced by guanines, resulted in white plants [*Nt*-IM13 (AAG+1); Figure 5B], suggesting complete loss of ATP synthase (Hager, 2002; Schmitz-Linneweber et al., 2005). Strongly reduced translation efficiency of these constructs was also observed with the dual-luciferase assay in *E. coli* (Figure 7). These findings indicate that frameshift compensation in *atpB* requires a homopolymeric tract. It was shown in bacteria that mRNAs encoding iterated lysine codons (either AAA or AAG) differ remarkably in efficiency and accuracy of protein synthesis. The presence of repetitive AAA codons reduced protein expression compared to the presence of synonymous AAG codons. This is because 70S ribosomes read AAA and AAG codons differently. Ribosomes that encounter multiple AAA codons undergo sliding, whereas no sliding occurs on AAG codons (Koutmou et al., 2015). These findings are in line with our observation that, in the dual-luciferase assay, an enhanced translation efficiency was detected in pEK4-*Nt*-IM11 (AAG) compared to the wild-type sequence that carries the oligoA stretch [pEK4-*Nt*-WT] (Figure 7).

Interestingly, the −2A deletion [*Nt*-IM18 (−2A)] results in an albino phenotype with rare green spots (Figure 8B, Figure 6C). *Nt*-IM18 (−2A) plants are phenotypically more similar to *Nt*-IM13 (AAG+1), and no AtpB peptides were detected in *Nt*-IM18 (−2A). This indicates that −2 or +1 ribosomal frameshifting are not active in chloroplasts (Table 1), and an A_8_ homopolymeric tract is also insufficient to effectively induce transcriptional slippage. This is not entirely unexpected in that the propensity of RNA polymerase to stutter is known to be strongly correlated with the length of the (homopolymeric) slippery site (Lin et al., 2015).

### What mechanism of recoding is acting in I-iota and transplastomic tobacco lines?

It can be challenging to distinguish between transcriptional slippage and PRF when monitoring it at a low level (Atkins et al., 2016). PRF often results in the formation of more than one protein. Likely due to instability of the truncated AtpB protein (Nishimura et al., 2017), our proteomic analyses detected only peptides corresponding to the corrected reading frame. However, two corrected protein variants (−1 and +2) were found for *Nt-*IM14 (+2A) (Figure 8F), lending circumstantial support to the operation of ribosomal frameshifting rather than RNA polymerase stuttering. Analysis of the predicted mRNA secondary structure revealed the presence of a stem-loop-type structure downstream the oligoA stretch that can serve as slip site for the ribosome (Figures 3C and 4B).

We have assessed the possibility of transcriptional slippage by cDNA analysis of the various mutants. No indication of transcriptional slippage for *Nt*-IM12 (+1A) and I-iota (+1A) (Figure 10A,B) was obtained, but spike-in experiments mixing of *Nt*-IM12 (+1A) or I-iota (+1) with the corresponding wild-type RNAs revealed a detection limit of approximately 15% (Supplemental Figure 2). Since the frame correction in *Nt*-IM12 (+1A) is estimated to be below 10% (Figure 5D), the operation of transcriptional slippage cannot be definitively excluded for *Nt*-IM12 (+1A).

Strikingly, in the transplastomic tobacco mutants, the +2A insertion into the oligoA tract is more efficiently compensated than the +1A insertion [*Nt*-IM14 (+2A) vs. *Nt*-IM12 (+1A), Figures 8B and 5B]. *Nt*-IM14 (+2A) plants are phenotypically most similar to wild type under mixotrophic conditions, and also can grow photoautotrophically and produce seeds (Figures 8B and 9A). Regarding their photosynthetic parameters, *Nt*-IM14 (+2A) plants display an intermediate phenotype between the wild type and *Nt*-IM12 (+1A) plants (Figures 8C and 9C, Table 3). In *Nt*-IM14 (+2A) plants, AtpB peptides with two or three lysines were detected, suggesting two modes of correction: +2 and −1 (Figure 8F). In other systems, several cases have been reported, where two different ribosomal frameshifts can occur at the same site (Caliskan et al., 2017; Li et al., 2019). As for *Nt*-IM12 (+1A), cDNA analysis revealed no indication of transcriptional slippage in *Nt*-IM14 (+2A) plants (Figure 10A). Importantly, in the cDNA sensitivity test, 5% of added RNA was detectable, when wild type and *Nt*-IM14 (+2A) RNAs were mixed (Figure 10C). Together with the absence of detectable RNA polymerase stuttering, this high sensitivity suggests that the correction occurs at the level of translation (i.e., by PRF). Work in microbial systems has revealed that the efficiency of alternative decoding can be highly variable, ranging from a few percent to up to 80% (Baranov et al., 2005; Caliskan et al., 2015). The frameshifting efficiency in *Nt*-IM14 (+2A) measured by the dual-luciferase reporter assay in *E. coli* was approximately 40% (Figure 7). The efficiency *in planta* may be lower, as evidenced by the measured ATP synthase activity in *Nt*-IM14 (+2A) plants, which was 18% of the activity in wild-type plants (Figures 8D and 9B).

### Functional significance of recoding in chloroplasts

In plastid genomes, homopolymeric nucleotide stretches are hotspots of mutations (Massouh et al., 2016). Homopolymeric tracts are selected against in bacterial genomes, but are retained more frequently in the genomes of endosymbionts. Transcriptional slippage has been proposed to play a critical role in rescuing gene function (Tamas et al., 2008; Wernegreen et al., 2010). Here polymerase stuttering at homopolymeric tracts can be considered as a mutation-compensating mechanism (Tamas et al., 2008). Whether or not transcriptional slippage occurs in chloroplasts and compensates frameshift mutations and/or generates additional transcript diversity, is currently unclear. If it occurs at the oligoA stretch near to 5’ end of *atpB*, its frequency is too low to be detected with the sensitivity of cDNA analysis (Figure 10, Supplemental Figure 2).

Programmed ribosomal frameshifting results in the formation of additional protein products, but can also correct indels at the level of translation (Ketteler, 2012; Atkins et al., 2016). Usually, ribosomal frameshifting is tightly regulated. Specific metabolites and aminoacyl-tRNA supply can affect the efficiency of ribosomal frameshifting (Ivanov et al., 2000; Caliskan et al., 2017). When we searched for other possible examples of alternative genetic decoding in the chloroplast, a sequence motif in the *cemA* (*ycf10*) gene (coding for an inner envelope protein likely involved in CO_2_ uptake) (Rolland et al., 1997) was identified. In contrast to *O. elata* (which was invested here), *O. glazioviana* and *O. parviflora* (*Oenothera* subgenus *Oenothera*) as well as *O. villaricae* and *O. picensis* (*Oenothera* subgenus *Munzia*) contain two ATG codons embedded in an oligoA stretch within the 5’ region of *cemA*. The oligoA stretches of the two pairs of species differ by an indel of 3 bp, and only the second ATG is in frame. To be used, the first ATG would require a +1 (+4) or −2 frameshift. Similar to *atpB*, the oligoA stretch could serve as slippery sequence motif for recoding (Greiner et al., 2008). To what extent recoding can result under certain experimental conditions even in gene fusion products, as suggested by the AtpB/AtpE fusion observed by Sears and Herrmann (1985) (also see Results), remains a matter of speculation.

Previous reports on the possibility of recoding in plants were based on bioinformatic prediction (Lin et al., 2015; van der Horst et al., 2019), *in vitro* analysis (Napthine et al., 2003) or indirect genetic evidence (Kohl and Bock, 2009). The data presented in this study provide clear genetic and biochemical evidence for the presence of recoding in the chloroplast *atpB* gene both in the spontaneous plastome mutant I-iota of *Oenothera* and a set of transplastomic tobacco plants. Its presence in chloroplasts may be a relic from the bacterial past of the chloroplast (Lin et al., 2015). To what extent the compensation of mutations by frameshifting is dependent upon the environmental conditions and/or the physiological status of the chloroplast, is an interesting question to be addressed in future studies.

## Material and Methods

### Plant material and growth conditions

*Oenothera* (evening primrose) wild type (*Oe*-WT; *O. elata* ssp. *hookeri* strain johansen Standard equipped with the chloroplast genome of *O. elata* ssp. *hookeri* strain hookeri de Vries; johansen Standard I-hookdV) and the I-iota mutant (johansen Standard I-iota) were previously described in Massouh et al. (2016) and cultivated according to Greiner and Köhl (2014). In *Oenothera,* photosynthetically incompetent chloroplast mutants can be kept as variegated plants, containing a mixed population of green nursery chloroplasts and mutated plastids. These plants are viable in soil because the green tissue feeds the non-photosynthetic mutant leaf tissue (Figure 1A). The combination of two chloroplast types in one plant is achieved by taking advantage of biparental chloroplast transmission, which is the rule in evening primroses. To this end, a maintainer strain containing green nursery plastids is crossed to the chloroplast mutant. This results in variegated offspring because vegetative sorting (sorting-out) of the two chloroplast types leads to tissues homoplasmic for either green or mutant chloroplasts in the same plant. Biparental transmission of plastids in *Oenothera* displays, however, maternal dominance. Consequently, the F1 generation offspring is usually either homoplasmic for the maternal plastome or heteroplasmic for the paternal and maternal plastomes. To increase the frequency of variegated (heteroplasmic) offspring in F1, a johansen Standard line was equipped with the slowly multiplying plastome IV of *O. parviflora* strain atrovirens Standard (IV-atroSt) as green nursery plastome (cf. Sobanski et al., 2019). This line was then used as parent in crosses with the faster multiplying I-iota plastome. Resulting F1 generations display up to 100% variegation for green and mutated plastomes in the progeny. Homoplasmic mutant tissue generated this way was used for molecular and physiological analyses in this work. For details on the genetics, see (Kutzelnigg and Stubbe, 1974; Kirk and Telney-Basset, 1978; Stubbe and Herrmann, 1982; Greiner, 2012; Sobanski et al., 2019).

For plant growth, *Oenothera* seeds were soaked in water for 12-16 h at 4°C, germinated on wet filter paper at 27°C in the light, followed by seedling transfer to soil. About one month after germination (early rosette stage) plants were vernalized at 4°C for seven days under a 10 h light/14 h dark regime. Plants were then returned to the greenhouse and cultivated under long-day conditions (16 h light, 180-220 µE m^-2^ s^-1^, 22 °C/8 h dark, 20 °C, humidity 50%).

Homoplasmic I-iota plants and wild-type control plants were cultivated in Magenta vessels (Sigma-Aldrich) under aseptic conditions (sterile culture). Seeds were sterilized following a vapor-phase sterilization protocol. Seeds in open microcentrifuge tubes were placed in a desiccator jar together with a glass beaker containing 100 mL of 13% [v/v] NaClO. Prior to sealing the desiccator jar, 3 mL concentrated HCl was carefully added to the bleach. Seeds were then sterilized in developing chlorine fumes for 5 h, followed by seeds ventilation in open tubes in a laminar flow hood for several hours to completely eliminate the chlorine gas. Subsequently, seeds were plated on germination medium containing ½ MS salts (Murashige and Skoog, 1962), 0.5% [w/v] Agargel (Sigma Aldrich), and 0.05% [v/v] Plant Preservative Mixture (Plant Cell Technology), adjusted to pH 5.8 with KOH, and incubated for 27°C in the light. After one week, seedlings were transferred to ½ MS medium with Agargel (see above) supplemented with 1.75% [w/v] glucose and cultivated at 16 h light 50 µE m^-2^ s^-1^/8 h dark and 24 °C.

*Nicotiana tabacum* cv. Petit Havana (*Nt*-WT) and *atpB*-*aadA* control plants (Rott et al., 2011) were raised from seeds in soil under controlled growth conditions (16 h light, 150-180 µE m^-2^ s^-1^, 25 °C/8 h dark, 20°C; humidity 60%). Transplastomic tobacco lines were cultivated in Magenta boxes on agar-solidified MS medium containing 3% [w/v] sucrose and supplemented with 500 mg/L spectinomycin under 16 h light, 50 µmol m^-2^s^-1^/8 h dark, 24 °C. For homoplasmy tests (Svab and Maliga, 1993; Bock, 2001), seeds were surface-sterilized by soaking in 70% [v/v] ethanol for two minutes, followed by soaking in 3% [v/v] NaClO containing a drop of Triton X-100 for 15 minutes. The seeds were then rinsed with sterilized water 5-7 times and plated on MS-based selective media containing 500 mg/L spectinomycin, or 500 mg/L streptomycin (Rott et al., 2011). Homoplasmic *Nt*-IM13 (AAG+1) and *Nt*-IM18 (−2A) seeds were obtained from stable *Nt*-IM13 (AAG+1)/WT and *Nt*-IM18 (−2A)/WT periclinal chimeras (plants with white leaf border carrying mutated chloroplast and green leaf blade with wild-type chloroplasts; cf. Greiner 2012), respectively.

### Test for homoplasmic I-iota tissue

Homoplasy of leaf sectors of I-iota was assessed using a sequence length polymorphism in the *clpP* gene that distinguishes the nursery plastome IV-atroSt (GenBank accession number EU262891.2) from the I-iota plastome that derives from I-hookdV (GenBank accession numbers KT881170.1; Massouh et al. 2016). The marker was amplified by standard PCR, employing the primer pair clpP_IvsIV_for and clpP_IvsIV_rev (Supplemental Table 4; cf. Figure 1B).

### Generation of transplastomic tobacco lines

Point mutations in *atpB* were introduced by site-directed mutagenesis. 177 bp from the 5’-part of *atpB* were cloned into the pEXA vector (Eurofins) and used as a template. Primer pairs used for PCR are listed in Supplemental Table 4. Reactions were carried out employing Phusion High Fidelity DNA polymerase according to the manufacturer’s instructions (Thermo Scientific). Non-mutated parental DNA was digested with the restriction enzyme DpnI for 1 h at 37°C. DpnI-treated DNA was transformed into DH5-alpha chemically competent cells and plated on LB medium supplemented with 100 µg mL^-1^ ampicillin. Single colonies were used for inoculation of overnight culture and plasmids isolated with the help of the QIAprep Spin Miniprep Kit (Qiagen). The presence of the mutation was verified by Sanger sequencing (LGC genomics). Positive clones were digested by CpoI and BspOI (Thermo Scientific) and the released *atpB* fragment was ligated into the similarly digested *aadA-atpB* plasmid (Figure 6A; Rott et al., 2011) using T4 ligase (Invitrogen).

Transplastomic lines were generated by biolistic transformation (Rott et al., 2011). Several homoplasmic lines were identified after the first regeneration round. Homoplasmy of the lines was verified by restriction fragment length polymorphism (RFLP) analysis (Figure 6B). Point mutations were confirmed by Sanger sequencing (LGC genomics).

### DNA gel blot and RFLP analyses

Total plant DNA was isolated by a rapid cetyltrimethylammonium bromide-based miniprep method (Doyle, 1990). DNA samples were digested with PvuII and NcoI, separated in 1% [w/v] agarose gels and blotted onto Amersham Hybond-XL membranes (GE Healthcare) according to the supplier’s recommendations. Specific probes were generated by PCR amplification (Figure 6A, primers are listed in Supplemental Table 4). For hybridization, α [^32^P] dCTP-labeled probes were generated by random priming (Megaprime DNA labeling kit; GE Healthcare). Hybridization was performed in Church Buffer (1% [w/v] BSA, 1 mM EDTA, 7% [w/v] SDS, 0.5 M Na_2_HPO4/H_3_PO_4_, pH 7.2) at 65°C overnight. Subsequently, the membranes were briefly rinsed with Wash Solution I (0.1% [w/v] SDS, 300 mM NaCl, 30 mM sodium citrate), then incubated under continuous agitation in Wash Solution I at room temperature for 20 min, and finally washed twice with Wash Solution II (0.1% [w/v] SDS, 75 mM NaCl, 7.5 mM sodium citrate) for 15 min at 65 °C. For signal detection, the membranes were incubated in phosphorimager cassettes, and the hybridization signals were visualized by scanning in aTyphoon TRIO+ Variable Mode Imager (Amersham Biosciences).

### Analysis of cDNA

RNA from *Oenothera* and tobacco leaves was extracted with the Spectrum Plant Total RNA kit (Sigma Aldrich). Genomic DNA was eliminated with the Turbo DNA-free kit (Ambion). cDNA was generated with the AccuScript High Fidelity cDNA kit (Agilent) according to the supplier’s recommendation and using gene-specific primers (Supplemental Table 4). Phusion High-Fidelity DNA polymerase (ThermoFisher Scientific) was used for subsequent PCR amplification.

### Prediction of RNA secondary structure

mRNA secondary structures were predicted with the mfold web server (Zuker, 2003).

### Photosynthesis measurements

Light-response curves of chlorophyll-*a* fluorescence parameters were determined at 22°C using the fiberoptics version of the Dual-PAM instrument (Heinz Walz GmbH). Linear electron flux and the redox state of the PSII acceptor side (qL) (Kramer et al., 2004) were measured after 30 min of dark adaptation. Light intensity was stepwise increased from 0 to 2000 μE m^−2^ s^−1^, with a measuring time for each light intensity of 150 s under light-limited conditions and 60 s under light-saturated conditions. After the measurement, the chlorophyll content and chlorophyll a/b ratio of the measured leaf segment were determined in 80% (v/v) acetone according to (Porra et al., 1989).

Proton conductivity of the thylakoid membrane (gH^+^) was used as a measure of ATP synthase activity. It was determined on intact leaves from the decay kinetics of the electrochromic shift signal (ECS) during a short interval of darkness. The experiments were performed at 22°C. Leaves were pre-illuminated for 5 min with saturating light (1000 μE m^−2^ s^−1^) so that photosynthesis was fully activated and ATP synthase activity was not limited by ATP consumption by the Calvin–Benson-Bassham cycle. The saturating illumination was interrupted by 15 s intervals of darkness, and the rapid first phase of the decay kinetic of the electrochromic shift was fitted with a single exponential decay function. Depending on the mutants measured, this fit was restricted to time intervals between 200 ms [for the wild type *Nt*-WT, *aadA-atpB* and *Nt*-IM11 (AAG)] and a maximum of 800 ms after the end of the actinic illumination [in case of mutants with very low ATP synthase activity such as *Nt*-IM12 (+1A), *Nt*-IM14 (+2A), or *Nt*-IM16 (−1A)]. The reciprocal value of the time constant was used as a measure of ATP synthase activity. Signals were measured and deconvoluted using a KLAS-100 spectrophotometer (Heinz Walz GmbH) as previously described (Rott et al., 2011).

### Thylakoid isolation

Leaves from 6-8 week-old *Oenothera* plants were harvested in the beginning of the light phase and shortly placed in ice water. 10 g of leave material was mixed with 200 mL Isolation Medium [50 mM HEPES/KOH (pH 7.6), 330 mM sorbitol, 2 mM EDTA, 1 mM MgCl_2_, 5 mM sodium ascorbate] and blended in a Waring blender (5 times 5 s each). Then, the homogenate was filtered through a double layer of mull and Miracloth (Merck), and the filtrate centrifuged at 5.000 g for 5 min at 4°C. Subsequently, the pellet was resuspended with a brush in 40 mL Wash Medium [50 mM HEPES/KOH (pH 7.6), 5 mM sorbitol)], homogenized in a 30 mL potter homogenizer (PYREX) until the chloroplast pellet was dissolved, and centrifuged at 5.000 *g*, 5 min, 4°C. The pellet was then again resuspended, homogenized in a potter homogenizer and centrifuged two times as above. Finally, the pellet was resuspended in 5 mL washing medium, layered on top of 85% Percoll solution [85% PBF-Percoll stock solution contains 3% [w/v] PEG 6000, 1% [w/v] bovine serum albumin, 1% [w/v] Ficoll 400; 330 mM sorbitol, 50 mM HEPEs/KOH (pH7.6), 2 mM EDTA, 1 mM MgCl_2_] and centrifuged at 10.000 *g* for 5 min at 4°C. Thylakoids were washed once with Wash Medium (5.000 g, 5 min, 4°C) and resuspended in Storage Buffer [50 mM HEPES/KOH (pH 7.6), 10% [v/v] glycerol, 5 mM MgCl_2_, 100 mM sorbitol], frozen in liquid N_2_ and stored at −80°C until use.

### Protein isolation and SDS-PAGE

Thylakoids membranes (20 µg chlorophyll equivalent) were pelleted by centrifugation (20,000 *g*, 10 min, 4 °C) and resuspended in 30 µL extraction buffer [25 mM BisTris/HCl (pH 7.0), 20% [v/v] glycerol, 0.25 g/l cOmplete protease inhibitor (Roche)]. Plant total proteins were extracted from 20 mg of frozen ground leaf material homogenized in 100 µL of extraction buffer. Membranes were solubilized by addition of 1% [w/v] n-dodecyl-β-D-maltoside (Sigma Aldrich) in the darkness on ice for 5 min. Finally, samples were denatured in Sample Buffer (62 mM Tris/HCl, pH 6.8, 10% [v/v] glycerol, 2% [v/v] sodium dodecyl sulfate) for 3 min at 95°C. SDS polyacrylamide gel electrophoresis with 9% [T] separation gels was performed according to (Laemmli, 1970). After electrophoresis, proteins were visualized by Coomassie staining. Separation gels were incubated in staining solution (0.1% [w/v] Coomassie Blue G, 0.1% [w/v] Coomassie Blue R, 45% [v/v] methanol, 9% [v/v] acetic acid) for 10 min under continuous agitation at room temperature. Gels were destained by incubation in 30% [v/v] methanol, 5% [v/v] acetic acid solution.

### Western blot analyses

Proteins separated by SDS-PAGE were transferred to polyvinylidene difluoride membranes (0.2 µm, Thermo Scientific) using a semi-dry electroblotting system (PeqLab). Primary antibodies produced in rabbit against AtpB and AtpE were purchased from Agrisera. As secondary antibody, an anti-rabbit IgG peroxidase conjugate was used (Agrisera). Immunochemical detection was performed using the ECL system (GE Healthcare) according to the supplier’s recommendation. Membranes were exposed to an X-ray film (Amersham TM ECL, GE Healthcare). Films were developed using an X-ray film developer (Hyper processor, GE Healthcare).

### Samples preparation for LC-MS/MS

For in-solution digestion, thylakoids from *Oe*-WT and I-iota (50 µg each) were solubilized in UTU buffer (6 M urea/2 M thiourea, pH 8.0) and subsequently incubated in reduction buffer (1 µg/µL DTT in ddH_2_O) for 1 h at 50°C followed by incubation with alkylation buffer (55 mM iodoacetamide in ddH_2_O) in the dark for 30 min at room temperature. The samples were then treated with endoproteinase Lys-C (0.5 µg/µL 10 mM Tris/HCl, pH 8.0) for four hours at room temperature and digested in 0.4 µg/µL of trypsin (both purchased from Promega) overnight at room temperature. Bovine serum albumin (BSA) was used as a control and treated identically. Digestion was stopped by addition of TFA (0.2% [v/v] final concentration). Peptides were purified using Zip tip C18 pipette tips (Millipore) according to the manufacturer’s instructions.

For in-gel digestion, SDS–PAGE-separated and Coomassie-stained protein bands were excised from the gel and destained using a mixture of 40% [v/v] acetonitrile and 60% 50 mM [v/v] NH_4_HCO_3_ at 37°C under continuous agitation. Gel pieces were dried by vacuum centrifugation and incubated with trypsin (modified sequencing grade, Roche) in 50 mM NH_4_HCO_3_ or Glu-C (sequencing grade, Promega) in 50 mM Tris/PO_4_, pH 7.8 for 12-16 h at 37°C. Proteolytic peptides were extracted by repetitive incubation with acetonitrile, 5% [v/v] formic acid, and again acetonitrile. The combined supernatants were dried by vacuum centrifugation, and peptides from were purified using Zip tip C18 pipette tips (Millipore) according to the manufacturer’s instructions.

### LC-MS/MS

Samples generated by in-solution digestion for quantification of AtpA, AtpB, and AtpE in *Oe*-WT and I-iota were subjected LC-MS/MS with free labeling approach. The peptides were resuspended in 2% (v/v) acetonitrile, 0.1 % (v/v) TFA and injected using nanoflow HPLC (Proxeon Biosystems) into the analytical column (length: 12 cm, diameter: 75 μm) filled with C18 (Reprosil C18; Dr. Maisch GmbH). Peptides were eluted from the column in a 90 minute linear gradient of 5 to 80% acetonitrile at a flow rate of 250 µL/min. Samples were analyzed using the selected reaction monitoring method (SRM). Fragmentation and detection were performed with a TSQ Quantum UltraTM Triple Stage Quadrupole Mass Spectrometer (Thermo Fisher Scientific). Specific peptide sequences unique for the target proteins (AtpA, AtpB, AtpE) were created using the Pinpoint software (version 1.0, Thermo Scientific) and are listed in Supplemental Table 3. Peptides obtained from mass spectrometry were analyzed with the Pinpoint software (Thermo Scientific). The proteomic datasets have been deposited in the ProteomeXchange Consortium (http://www.proteomexchange.org) via the PRIDE partner repository with the dataset identifier PXD020246.

Samples generated by in-gel digestion were resuspended in 5% (v/v) acetonitrile and 0.1% (v/v) formic acid. Peptides were separated on a C18 reversed-phase analytical column (Acclaim PepMap100, Thermo Fisher Scientific) using an Easy-nLC 1000 liquid chromatography system (Thermo Fisher Scientific). Peptides were eluted in a 28-min-long non-linear 5% - 34% acetonitrile gradient in 0.1% formic acid and 5% DMSO at a flow of 300 nL min^-1^. Subsequently, the column was cleaned for 10 min with 85% acetonitrile in 0.1% formic acid and 5% DMSO. Eluted peptides were transferred to an NSI source and sprayed into an Orbitrap Q-Exactive Plus mass spectrometer (Thermo Fisher Scientific). The MS was run in positive ion mode. For full MS scans, the following settings were used: resolution: 70000, AGC target: 3E6, maximum injection time: 100 ms, and scan range: 200 to 2000 m/z. For dd-MS^2^, the following settings were used: resolution: 175000, AGC target: 1E5, maximum injection time: 50 ms, loop count: 15, isolation window: 4.0 m/z, and NCE: 30. In addition, the following data-dependent settings were used: underfill ratio: 1%, apex trigger: off, charge exclusion: unassigned, 1, 5, 5–8, >8, peptide match: preferred, exclude isotypes: on, and dynamic exclusion: 20.0 s.

The raw files obtained from Xcalibur (Thermo Fisher Scientific) were converted into mgf files using MSConvert from the Proteowizard portal. The files were analyzed by the Mascot server with an in-house database containing protein sequences from several plant species POTbase MS (Moreno et al., 2018), or a database containing the AtpB sequence and the potential variations in lysine content (summarized in Table 1, for sequences see Supplemental Table 1).

### Multiple sequence alignments

ClustalW (Thompson et al., 1994) implemented in the MegAlignPro software (DNAStar) was used to perform multiple sequence alignments. DNA sequences were obtained from NCBI Gene (https://www.ncbi.nlm.nih.gov/gene/). Sequences of *atpB* with the following gene ID were used to generate the alignment: 844757 (*Arabidopsis thaliana*), 29141377 (*Oryza sativa*), 4099984 (*Solanum tuberosum*), 845170 (*Zea mays*), 802790 (*Oenothera elata*), and 800515 (*Nicotiana tabacum*).

### Dual-luciferase reporter assay

The dual luciferase reporter plasmid was designed on the basis of the pEK4 vector (Kramer and Farabaugh, 2007). 150 bp from the 5’-region of *atpB* from *N. tabacum* and *O. elata* (GenBank accession numbers Z00044.2 and KT881170.1, respectively) containing the oligoA stretch were cloned into the polylinker of the insertion window of pEK4 using BglII and BamHI restriction sites. Final constructs were transformed into NEB Express chemically competent *E. coli* cells (C3037I, NEB). Cultures of three individual clones were grown at 37°C in LB medium supplemented with 100 µg mL^-1^ of ampicillin. Upon reaching an OD of 0.4, equal volumes of suspension were collected. Crude extracts were prepared according to (Meydan et al., 2017). The activities of firefly and Renilla luciferases were measured using the Dual-Luciferase Reporter Assay System (Promega). Luminescence was followed in 96-well plates in a ClarioStar micro plate reader (BMG Labtech). The PRF efficiency was calculated according to (Grentzmann et al., 1998).

### Transmission electron microscopy

*Oenothera* leaf pieces of 2 mm^2^ were fixed with 2.5% [w/v] glutaraldehyde and 2% [w/v] paraformaldehyde in 0.2 M sodium cacodylate buffer (pH 7.4) for at least 4 h at 4°C. Samples were then post-fixed in 1% [w/v] OsO_4_ at 4°C overnight, stained in 2% [w/v] aqueous uranyl acetate for 2 h and dehydrated gradually in 30, 50, 70, 80, 90 and 100% [v/v] ethanol followed by washing in 100% [v/v] propylene oxide two times. The samples were then embedded in low-melting Spurr epoxy resin (Agar Scientific), degassed and cured at 65 °C for 24 h. Thin sections (60-70 nm) were obtained with a Leica Ultracut UC 6 (Leica Microsystems), mounted on 150 mesh nickel grids, counterstained with uranyl acetate followed by lead citrate, and examined with a transmission electron microscope at 120 kV (EFTEM, Zeiss).

## Acknowledgments

The authors thank Stephan Obst for help with plant transformation, Wolfram Thiele for help with photosynthesis analyses and ATP synthase activity measurements, Eugenia Maximova for transmission electron microscopy, David Rolo (both MPI-MP) for assistance with MS analysis, and Barbara B. Sears (MSU, USA) and Reinhold Herrmann (LMU, Germany) for critical reading and valuable comments on the manuscript. We are grateful to Philip J. Farabaugh (UMBC, USA) for providing the pEK4 plasmid, and to Britta Hausmann and Florian Hundert (MPI-MP Green Team) for plant cultivation. This work was supported by a grant from the Deutsche Forschungsgemeinschaft (GR 4193/1-2) to S.G. and by the Max Planck Society to S.G, M.A.S., and R.B.

## Author contribution

I.M., A.Z., A.M. performed protein analysis (western blot and samples preparation for MS). E.M. and W.S. performed MS analysis. I.M., M.R, L.Y.R., and S.R. generated and identified tobacco transplastomic lines. M.A.S. performed photosynthesis analysis and measured the activity of the ATP synthase. All authors analyzed and discussed the data. I.M. and S.G. designed the study. I.M. wrote the manuscript with significant contribution from S.G., M.A.S and R.B. All authors approved the final version.

